# Sirtuin 1 Is Required for Optimal Mammarenavirus Multiplication

**DOI:** 10.64898/2026.07.23.740332

**Authors:** Haydar Witwit, Roaa Khafaji, Patricia Mingo-Casas, Ana-Belén Blázquez, Miguel A. Martín-Acebes, Juan Carlos de la Torre

**Affiliations:** Department of Immunology and Microbiology, The Scripps Research Institute, La Jolla, CA 92037, USA; Independent researcher, Chula Vista, CA, 91910, USA; Department of Biotechnology, Instituto Nacional de Investigación y Tecnología Agraria y Alimentaria, Consejo Superior de Investigaciones Científicas (INIA-CSIC), 28040, Madrid, Spain; Universidad Autónoma de Madrid (UAM, Escuela de Doctorado), Madrid, Spain

**Keywords:** mammarenaviruses, LCMV, hemorrhagic fever, sirtuin, SIRT1, neutral sphingomyelinase 2, nSMase2, host-directed antiviral, viral life cycle

## Abstract

Mammarenaviruses (MaAv) cause persistent infections in diverse rodent reservoirs worldwide and several are zoonotic pathogens with an important public-health burden in their endemic regions. Moreover, the globally distributed MaAv lymphocytic choriomeningitis virus (LCMV) is an underrecognized pathogen of clinical significance in congenital infections and immunocompromised individuals. The lack of FDA-approved vaccines or antivirals for MaAv infections underscores the urgent need for novel anti-MaAv therapeutic strategies. Neutral sphingomyelinase 2 (nSMase2) was recently identified as a host factor contributing to LCMV multiplication, and its inhibitor cambinol exhibits dose-dependent antiviral activity against LCMV but the underlying mechanisms remain undefined. Here, we show that cambinol disrupts multiple stages of the LCMV life cycle. Cambinol inhibits the pH-dependent fusion event mediated by MaAv glycoprotein, a step required for completion of virus cell entry. It also reduces viral ribonucleoprotein (vRNP)-directed genome replication and transcription and impairs the budding activity of the virus matrix Z protein. Cambinol also inhibits sirtuins 1 and 2 (Sirt-1 and Sirt-2), two NAD^+^-dependent protein deacetylases with pleiotropic roles in cellular metabolism and stress responses, raising the question of whether cambinol anti-LCMV activity reflects nSMase2 inhibition alone or also involves sirtuin-dependent pathways. LCMV multiplication was significantly reduced in SIRT1, but not SIRT2, knockout (KO) cells, uncovering a pro-viral role for Sirt-1 in the LCMV life cycle. Consistent with this finding, LCMV vRNP activity and production of infectious progeny were reduced in SIRT1 KO cells. These findings identify Sirt-1 as a host factor required for optimal LCMV multiplication. Sirt-1 inhibitors are in clinical development for oncological and neurological indications, raising the possibility of repurposing Sirt-1 inhibitors as host-directed antivirals (HDAs) against human pathogenic MaAv.

**Abstract figure:** 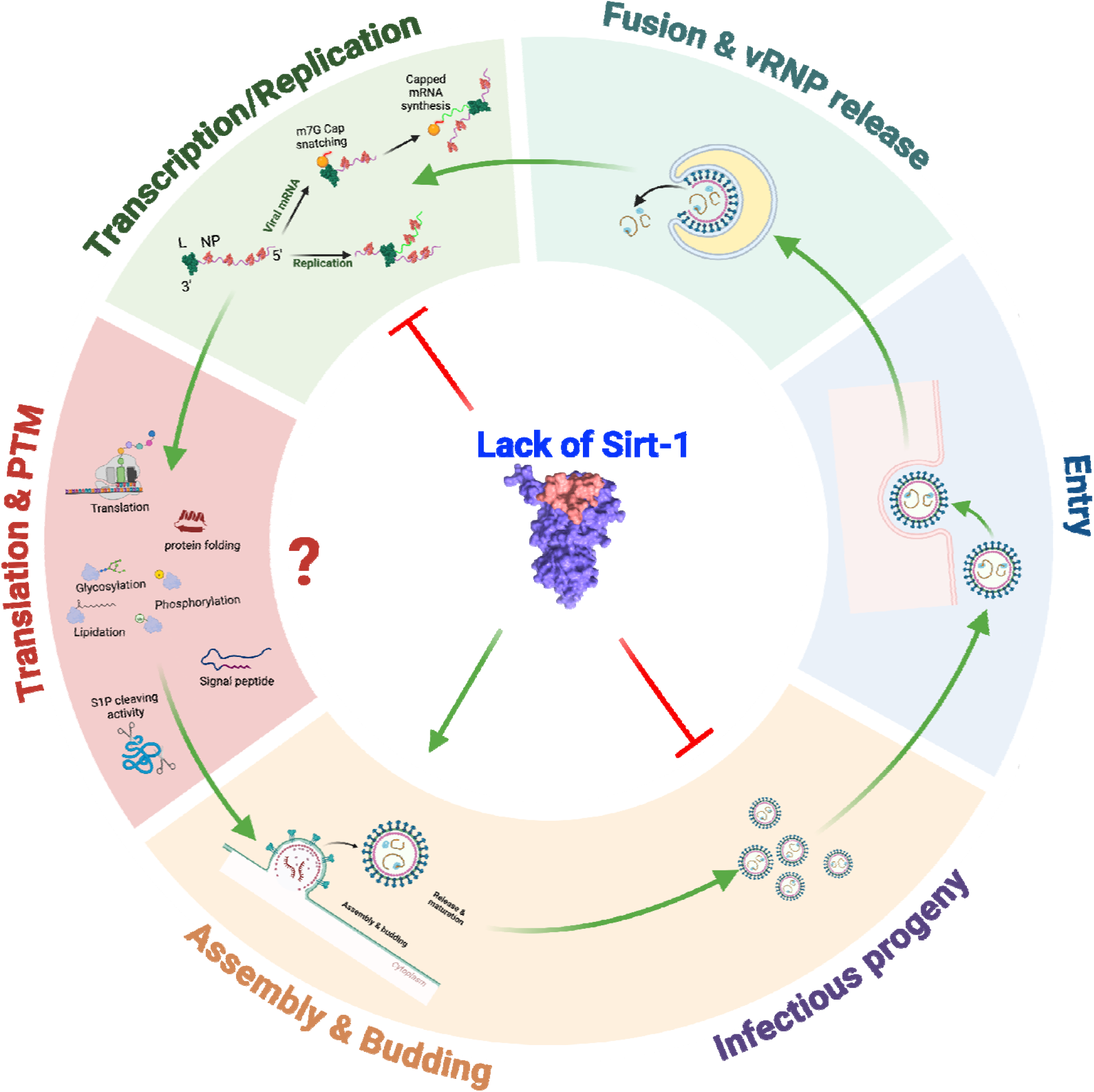

## 1. Introduction

Mammarenaviruses (MaAv) cause chronic infections in their rodent reservoirs worldwide, with human infections occurring primarily through mucosal exposure to aerosolized infectious material or direct contact with contaminated material [1]. Several MaAv cause hemorrhagic fever (HF) diseases in humans and represent important public health problems in their endemic regions. The Old World MaAv Lassa virus (LASV), endemic to Western Africa, is estimated to infect over 500,000 individuals annually [2], resulting in a high number of Lassa fever (LF) cases, an HF associated with high morbidity and case fatality rate as high as 69% among hospitalized confirmed LF cases [1,3,4]. Expanding endemic regions, climate-driven ecological changes, population movement, and potential for zoonotic spillover have raised concerns that LASV incidence and geographic distribution will continue to increase [5,6]. The New World MaAv Junin virus (JUNV), the causative agent of Argentine HF, and several other New World MaAv cause severe diseases throughout South America [7,8]. Additionally, the globally distributed prototypic MaAv lymphocytic choriomeningitis virus (LCMV) is an underrecognized human pathogen of clinical significance, particularly in congenital infections and in immunocompromised individuals [9,10]. Moreover, the significant seroprevalence of LCMV within different populations across the world, including the United States, has raised the question of whether LCMV may contribute to the many cases of undiagnosed aseptic meningitis reported yearly [11,12].

Despite their significant impact on human health, there are currently no FDA-approved MaAv vaccines or antivirals. Current treatment for MaAv infections is limited to the off-label use of ribavirin, whose therapeutic efficacy remains controversial [13]. Several direct-acting antivirals (DAAs) are under investigation, including the polymerase inhibitors favipiravir and 4’-fluorouridine and the entry inhibitors LHF-535 and ARN-75039, as well as antibody-based therapies [14–20]. However, DAA development can be complicated by toxicity and resistance emergence, driven in part by the high genetic diversity and mutational plasticity of MaAv RNA genomes [21–25]. Antibody-based therapies, while promising in preclinical models, face significant challenges including high cost, cold chain distribution requirements, and parenteral administration, which may restrict their use to specialized treatment centers [18]. Host-directed antivirals (HDAs), which target host factors and cellular processes hijacked by viruses for their multiplication, offer complementary advantages: because HDAs act on host rather than viral targets, they impose a higher genetic barrier to resistance emergence and have the potential to act as broad-spectrum antivirals against multiple members of a virus family that share common host dependencies [26,27].

As with other enveloped viruses, lipid metabolism plays an important role in MaAv multiplication. Specifically, LCMV infection induces extensive remodeling of the host cell lipid landscape, with particularly prominent alterations in sphingolipid and fatty acid metabolic pathways [28]. Among the sphingolipid-related host factors implicated in viral replication, neutral sphingomyelinase 2 (nSMase2) has attracted particular attention [27–30]. nSMase2, encoded by the SMPD3 gene, is the primary sphingomyelinase isoform responsible for the hydrolysis of sphingomyelin to ceramide at the cytoplasmic leaflet of the plasma membrane and at the Golgi apparatus, and plays central roles in membrane organization, exosome biogenesis, stress signaling, and autophagy [28,31–34]. A growing body of evidence implicates nSMase2 as a pro-viral host factor across diverse virus families. In HIV-1, nSMase2 interacts directly with the Gag polyprotein at plasma membrane assembly sites, where local ceramide generation drives membrane curvature required for virion budding and maturation. Accordingly, pharmacological or genetic inhibition of nSMase2 results in production of morphologically aberrant HIV-1 particles with impaired infectivity and substantially delays or eliminates viral rebound in humanized mouse models of HIV-1 infection [29,30]. Likewise, nSMase2 activity has been shown to support replication of West Nile and Zika viruses, and the selective nSMase2 inhibitor DPTIP exhibits potent antiviral activity against both flaviviruses [35]. nSMase2-dependent ceramide generation has also been linked to the formation of replication organelles during SARS-CoV-2 infection [36]. Consistent with these findings, both sphingomyelin synthase 1 (SMS1) and nSMase2 have been identified as pro-viral host factors in LCMV-infected cells [27,28].

Cambinol, a small-molecule inhibitor of nSMase2, exhibits dose-dependent antiviral activity against LCMV in cell culture [28]. However, cambinol also potently inhibits the NAD^+^-dependent deacetylases sirtuin 1 (Sirt-1) and sirtuin 2 (Sirt-2) [31], and therefore its anti-LCMV activity could be attributable to inhibition of either nSMase2 or sirtuins, or both. Sirtuins have pleiotropic roles in cellular metabolism, stress responses, transcriptional regulation, DNA repair, and cell cycle control [37]. Sirt-1 is predominantly nuclear but shuttles to the cytoplasm, and deacetylates both histone and non-histone substrates to regulate gene expression, autophagy, and the integrated stress response (ISR) [37]. Sirt-2 is primarily cytoplasmic and has established roles in cytoskeletal dynamics and mitotic regulation [37].

The roles of sirtuins in viral infection are broad and highly context dependent. On one hand, sirtuins can act as a viral restriction factor. Genetic perturbation and pharmacological inhibition of sirtuins have been shown to increase progeny production of both DNA and RNA viruses [38]. Thus, Sirt-1 restricts multiplication of enterovirus 71 (EV71) by directly binding the viral RNA-dependent RNA polymerase and attenuating its activity, as well as interacting with the 5’UTR of the EV71 RNA genome to inhibit viral replication and translation [39]. Likewise, Sirt-1 silences Kaposi’s sarcoma-associated herpesvirus (KSHV) lytic replication activator RTA at the chromatin level, maintaining viral latency, and Sirt-1 inhibition promotes lytic reactivation of KSHV [40]. On the other hand, Sirt-1 has been shown to be a positive regulator of hepatitis B virus (HBV) replication via targeting the transcription factor AP-1, and its inhibition suppresses HBV replication [41]. Whether sirtuins contribute to MaAv multiplication and through what mechanism has not been investigated.

In this study, we sought to dissect the relative contributions of nSMase2 and sirtuin inhibition to the anti-LCMV activity of cambinol. We found that cambinol inhibits the pH-dependent virus glycoprotein-mediated fusion activity, as well as vRNP-directed replication and transcription, and budding activity of the viral matrix Z protein. In addition, we show that LCMV multiplication is significantly reduced in SIRT1, but not SIRT2, knockout cells, uncovering a previously unrecognized pro-viral role for Sirt-1 in LCMV-infected cells. Mechanistic studies demonstrated that Sirt-1 absence impairs vRNP-directed replication and transcription of the viral genome and reduces infectious progeny production. These findings reveal an unexpected link between sirtuin-regulated cellular homeostasis and distinct stages of the MaAv life cycle, including viral RNA synthesis and particle release. Sirt-1 inhibitors are in clinical development for oncological and neurological indications [42], raising the possibility of repurposing them as host-directed antivirals (HDAs) against human pathogenic MaAv.

## 2. Materials and Methods

### 2.1. Cells and Viruses

All cell lines, including Human A549 (ATCC CCL-185), were cultured in Dulbecco’s Modified Eagle Medium (DMEM; ThermoFisher Scientific, Waltham, MA, USA) supplemented with 10% heat-inactivated fetal bovine serum (FBS), 2 mM L-glutamine, 100 μg/mL streptomycin, and 100 U/mL penicillin (Cat. No. 10378016, ThermoFisher Scientific, Waltham, MA, USA). Flp-In T-REx HEK293 or their isogenic mutants, sirtuin 1 or 2 knockout (SIRT1 or 2 KO) [43], cell lines were a gift from the Kwon lab (Department of Cellular Biology and Anatomy, Medical College of Georgia, Augusta University, Augusta, GA, USA). Recombinant viruses are as follows: rLCMV/GFP-P2A-NP (referred to as rLCMV/GFP), expressing green fluorescent protein (GFP) fused to the LCMV nucleoprotein (NP) via a P2A ribosomal skipping sequence; rLCMVΔGPC/ZsG-P2A-NP (rLCMVΔGPC/ZsG) [44], a single-cycle variant expressing Zoanthus sp. ZsG has been described previously. All the experiments with LCMV were conducted in the biosafety level 2 (BSL-2) laboratory at The Scripps Research Institute. Reverse genetics [45] was used to rescue rLCMV/mCherry-P2A-NP (referred to as rLCMV/mCherry), expressing mCherry protein (mCherry) fused to the LCMV nucleoprotein (NP) via a P2A ribosomal skipping sequence.

### 2.2. Antibodies and Compounds

Cambinol (Cat. No. HY-100732, MedChemExpress, NJ, USA) was dissolved in DMSO (Cat. No. D2650-100, Sigma-Aldrich (St. Louis, MO, USA)) at either 10 or 100 mM and stored in aliquots at ™20 °C. Entry inhibitor L862-0270 (also known as F3406) was obtained from ChemDiv, Inc. (Cat. No. L862-0270, San Diego, CA, 92130 USA). Ribavirin was purchased from Sigma-Aldrich (St. Louis, MO, USA) (Cat. No. R9644). Sodium citrate buffer (0.1 M, pH 5.0, sterile) was obtained from bioWorld (MFG No. 40121003-1, Dublin, OH, USA). The VL4 rat monoclonal antibody against NP was obtained from Bio X Cell (West Lebanon, NH, USA).

### 2.3. Cell Viability (CC_50_) and Viral Inhibition Half Maximal Effective Concentration (EC_50_)

Cell viability was assessed using the CellTiter 96 AQueous One Solution Reagent (G3580, Promega, Madison, WI, USA), which quantifies viable cells based on the conversion of a tetrazolium compound to a formazan product by NADPH or NADH in metabolically active cells. A549 cells were seeded in 96-well clear-bottom plates (4.0 × 10^4^ cells/well), treated with 2-fold serial dilutions of each compound, and incubated for 72 h. Following treatment, CellTiter 96 AQueous One Solution was added and incubated for 35 min at 37 °C in 5% CO_2_. Absorbance was measured using a Cytation 5 reader (BioTek, Agilent, Santa Clara, CA, USA), and values were normalized to vehicle (DMSO)-treated controls, set at 100%. The half-maximal cytotoxic concentration (CC_50_) was calculated using GraphPad Prism v11 (Prism11). Cell viability was also determined using DAPI staining. For determination of the half-maximal effective concentration (EC_50_), A549 cells were seeded in 96-well clear-bottom black plates (4.0 × 10^4^ cells/well), and 20 h later infected (MOI = 0.05) with rLCMV/GFP or rLCMV/mCherry. After 90 min of adsorption, the viral inoculum was removed and replaced with media containing test compounds. At 72 h post-infection, cells were fixed with 4% paraformaldehyde (PFA), and GFP or mCherry expression was measured by fluorescence using a Cytation 5 reader. Fluorescence readings were normalized to DMSO-treated controls (100%), and EC_50_ values were calculated using Prism11. The selectivity index (SI) for each compound was determined as the ratio of CC_50_ to EC_50_.

### 2.4. Viral Growth Kinetics

As described [24], cells were infected in 24-well plates at the indicated cell seeding density and MOI. After 90 min of adsorption at 37 °C and 5% CO_2_, the virus inoculum was removed, and fresh media with the indicated compounds and concentrations was added. At specified times post-infection, cell culture supernatants (CCS) were collected, and viral titers were determined by focus-forming assay (FFA).

### 2.5. Virus Titration

Virus titers were determined by focus-forming assay (FFA) as described [46]. Briefly, serial 10-fold dilutions of samples were prepared in DMEM containing 2% FBS and used to infect Vero E6 cell monolayers seeded in 96-well plates (3 × 10^4^ cells/well) and adsorbed for 2 h at 50 μL volume. Inocula were removed and replaced with 0.5% methyl cellulose (MC) containing media (2x EMEM, 2x MC, supplemented with 10% heat-inactivated fetal bovine serum (FBS), 2 mM L-glutamine, 100 μg/mL streptomycin, and 100 U/mL penicillin). At 24-48 h post-infection, cells were fixed with 4% paraformaldehyde in phosphate-buffered saline (PBS), and infection foci were visualized by epifluorescence based on GFP expression from rLCMV/GFP-infected cells using a fluorescence microscope.

### 2.6. LCMV Cell-Based Minigenome (MG) Assay

The LCMV cell-based MG assay was performed as described [24]. HEK293T single cell suspensions were prepared to obtain a total of 4.84 × 10^6^ cells in 5.2 mL of complete medium in a 50 mL conical tube. A transfection mixture (TM; 800 μL) was added to reach a final volume of 6 mL, and the suspension was incubated for 10 min at room temperature. Following incubation, 3.6 mL of complete medium was added and mixed gently by pipetting to ensure uniform distribution, resulting in a final TM volume of 9.6 mL. TM was added (80 μL/well) to poly-L-lysine-coated wells (96-well plate) pre-spotted with 20 μL of 5× cambinol. Wells were mixed twice with an electronic pipette, and plates were placed on a level surface platform for 15 min before being transferred to an incubator at 37 °C in a humidified atmosphere with 5% CO_2_.

The TM consisted of Lipofectamine 3000 (2.5 μL/μg DNA; Cat. No. L3000008, ThermoFisher Scientific, Waltham, MA) with Pol II-driven expression plasmids encoding T7 RNA polymerase (pC-T7, 1.32 μg), nucleoprotein (NP; pC-NP, 0.19 μg), and L polymerase (pC-L, 1.58 μg), along with a T7 promoter-driven plasmid expressing an LCMV S MG encoding the GFP reporter (pT7-MG/GFP, 1.32 μg) [24]. The LCMV MG transfection mixture was prepared by combining 403.75 μL of Opti-MEM containing plasmid DNA and 25 μL of P3000 reagent with 403.75 μL of Opti-MEM (Cat. No. 31985070, ThermoFisher Scientific, Waltham, MA, USA) containing 37.5 μL of Lipofectamine 3000, yielding a total volume of 806.6 μL. The mixture was incubated for 20 min at room temperature to allow for complex formation. At the indicated end timepoint, the cells were fixed with 4% paraformaldehyde (PFA), washed with PBS, then stained with DAPI. Images and GFP signal readings were collected using the Keyence BZ-X710 and Cytation 5 reader, respectively.

### 2.7. Time of Addition Assay

A549 cells were seeded in 96-well plates at a density of 6 × 10^4^ cells/well. The following day, cells were infected with the single-cycle infectious rLCMVΔGPC/ZsG (MOI = 0.5) and treated with cambinol (100 μM) or vehicle control (VC) either 1 h before infection (−1 h) or 1 h post-infection (+1 h). The LCMV entry inhibitor F3406 (10 μM) was included as a control treatment. At 48 hpi, ZsGreen-positive (ZsG^+^) cells were quantified using the Cytation 5 Reader (BioTek, Agilent, Santa Clara, CA, USA). Fluorescence values were normalized to those of VC-treated infected cells, and results represent the mean of three biological replicates.

### 2.8. GPC-Mediated Fusion Assay

HEK293T cells were seeded in poly-L-lysine-coated 24-well plates at a density of 2.5 × 10^5^ cells/well. The following day, cells were transfected with a pCAGGS plasmid encoding GFP (50 ng/well) along with either an empty pCAGGS vector or pCAGGS plasmids expressing LCMV or LASV GPC (1 μg/well) using Lipofectamine 3000, following the manufacturer’s instructions. After 5 h, the transfection medium was removed, cells were washed once, and fresh medium with or without cambinol was added. At 24 h post-transfection, cells were exposed to either acidified (pH 5.0) or neutral (pH 7.2) medium for 15 min, washed with DMEM, and returned to DMEM containing 10% FBS. Cells were then monitored over time for syncytia formation using fluorescence microscopy. Following observation, cells were fixed with 4% PFA, washed with PBS, and imaged at 20× magnification using a Keyence BZ-X710 microscope.

### 2.9. Z Budding Assay

Z budding activity was assessed using the described cell-based Z budding assay [47]. HEK293T cells were seeded in poly-L-lysine-coated 12-well plates at a density of 3.5 × 10^5^ cells/well. After overnight incubation, cells were transfected with 2 μg of either pC-LCMV-Z-Gaussia luciferase (GLuc), pC-LCMV-mutant Z [G2A]-GLuc, or pC-LASV-Z-GLuc using Lipofectamine 2000 (2.5 μL/μg DNA). Following a 5 h incubation, the transfection mixtures were replaced with fresh media containing the indicated compounds. At 48 h post-transfection, CCSs containing virion-like particles (VLPs) were collected and clarified by low-speed centrifugation to remove cell debris. Aliquots (20 μL) of each CCS sample were transferred to 96-well black plates (VWR, West Chester, PA, USA), followed by the addition of 50 μL Pierce Gaussia Luciferase glow assay kit reagent (Cat no. P16160, Thermo Fisher Scientific, Waltham, MA). Whole-cell lysates (WCLs) from the same wells were prepared to assess cell-associated GLuc activity. Luminescence was measured using the Cytation 5 Reader. Budding efficiency was calculated as CCS-GLuc/(CCS-GLuc + WCL-GLuc).

### 2.10. Statistical analysis

All statistical analyses were conducted, as indicated in the respective assays, using GraphPad Prism software v11 (GraphPad Software, Boston, MA, USA, www.graphpad.com). P value < 0.05, 0.01, 0.001, 0.0001 represented as *, **, ***, ****, while p value > 0.05 represented as ns (not significant). One-way or two-way ANOVA with multiple comparison correction with Tukey, Šidák or Dunnett methods were implemented when recommended.

## 3. Results

### 3.1. Dose-Dependent Effect of Cambinol on LCMV Multiplication

First, we generated an rLCMV/mCherry variant alongside the GFP-tagged strain to leverage the enhanced photostability and monomeric properties of mCherry, which improves signal quality and the potential use for multiplexing with other viruses or cells lines expressing different markers with different spectral profile. We characterized the rLCMV/mCherry in both multistep growth kinetic, MSGK in BHK21 and A549 cells (Figure 1A) and dose-response (DR) assay in A549 cells (Figure 1B, C). rLCMV/mCherry was comparable to rLCMV/GFP in the DR assay, and comparable to LCMV WT in thr MSGK. Cambinol exhibited a potent dose-dependent inhibitory effect on rLCMV/GFP and rLCMV/mCherry multiplication in A549 cells, with EC_50_ values of 35.66 and 35.32 μM, respectively, at 72 hpi (Figure 1B). The inhibitory effect of cambinol on LCMV multiplication was not a consequence of drug-induced cell toxicity, as cambinol exhibited a CC_50_ value > 200 μM (DAPI) and 138.3 μM (the CellTiter 96 AQueous One assay) and selectivity index (SI = CC_50_/EC_50_) of 3.9, a modest, yet clear separation (Figure 1B). While modest, this SI indicates that the observed antiviral activity is not attributable to cytotoxicity under the conditions tested. The difference between CC50 values obtained by the two viability assays (>200 μM by DAPI and 138.3 μM by CellTiter 96 AQueous One) likely reflects the distinct readouts of these methods: DAPI quantifies nuclear count as a measure of cell number, whereas CellTiter 96 AQueous One measures metabolic activity, which may be more sensitive to cytostatic effects at sub-lethal concentrations. The SI of 3.9 was calculated using the more conservative CC_50_ value of 138.3 μM.

**Figure 1.**
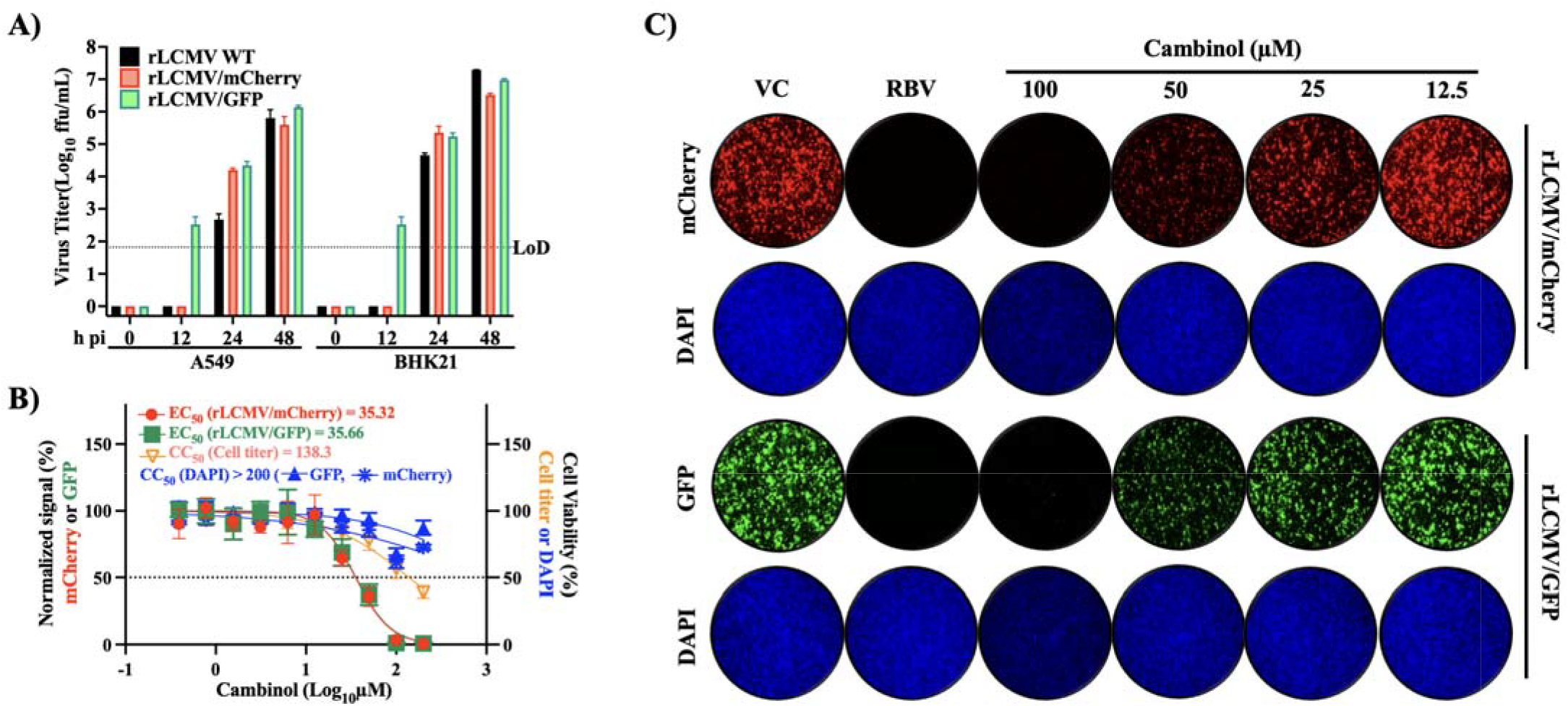
Cambinol exhibits a dose-dependent inhibitory effect on LCMV multiplication in human A549 cells. A) rLCMV/mCherry and rLCMV/GFP exhibit similar multi-step growth kinetics and peak titers in cultured cells. A549 or BHK21 cells were seeded at 5 × 10^5^ cells/well into a 24-well plate, infected with LCMV/WT (MOI = 0.01), rLCMV/GFP or rLCMV/mCherry (MOI = 0.05). At the indicated hours post-infection (h pi) virus titers in CCSs were determined in Vero E6 cells. B) A549 cells were seeded at 6 × 10^4^ cells/well into a 96-well plate, infected (MOI = 0.05) with rLCMV/GFP or rLCMV/mCherry and treated with cambinol at the indicated concentrations. At 72 hpi, cell viability was determined using the CellTiter 96 AQueous One assay, followed by fixation of cells in 4% PFA and DAPI staining to assess total cell number. GFP, mCherry and DAPI signals were quantified using Cytation 5 and normalized (%) to VC-treated infected controls. Results show the mean and SD of four biological replicates. EC_50_ and CC_50_ values were calculated using a variable slope (based on four parameters) model. C) Representative IF images of the dose-response assay shown in B. DAPI staining was used to visualize nuclei. Images were taken at 4× magnification using a Keyence BZ-X710 microscope. Ribavirin (RBV) (100 µM), a validated inhibitor of LCMV multiplication was used as a control.

### 3.2. Effect of Cambinol on LCMV Cell Entry

To assess whether cambinol affected a cell entry or post-cell entry step of LCMV infection we conducted a time of addition assay using a single-cycle infectious rLCMVΔGPC/ZsG to prevent the confounding effects of multiple rounds of replication without the need for NH4Cl treatment (Figure 2A). Treatment with cambinol (at 25, 50 and 100 μM) at both −1 h and +1 h of infection reduced the ZsG signal compared to VC treatment (Figure 2A). In contrast, treatment with the LCMV entry inhibitor L862-0270 led to a reduction in ZsG^+^ cells when administered at −1 h, but not at +1 h, of infection. LCMV cell entry is completed in less than 60 min [48]. Hence, results of the time of addition assay indicated that cambinol treatment was able to disrupt a post-cell entry step.

**Figure 2.**
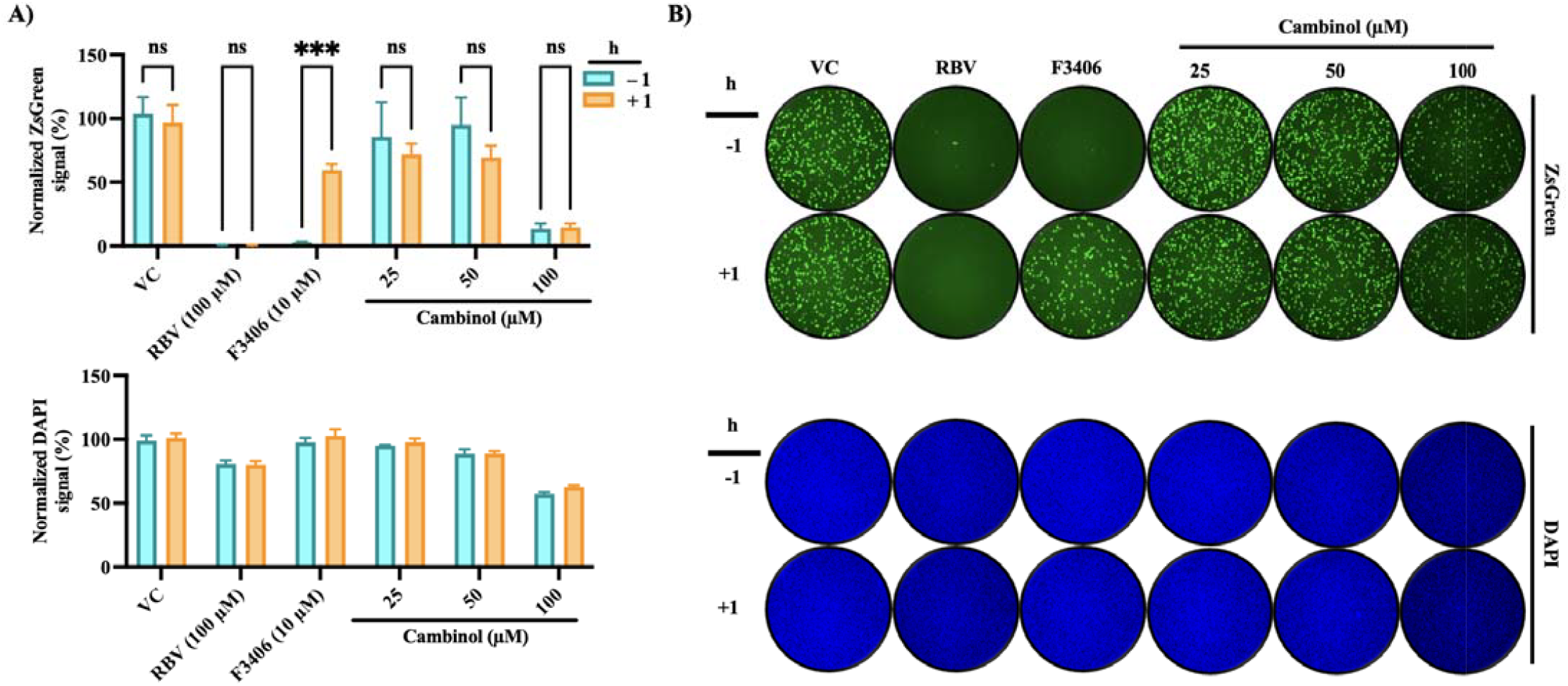
Effect of cambinol on LCMV cell entry. A) Time-of-addition assay. A549 cells were seeded in 96-well plates at 8 × 10^4^ cells/well. At 16 h post-seeding, cells were infected (MOI = 0.5) with a single-cycle infectious recombinant LCMV expressing ZsGreen (rLCMVΔGPC/ZsG). Cambinol (25, 50 or 100 µM) or VC was added either 1 h prior to infection (−1 h) or 1 h post-infection (+1 h). The LCMV entry inhibitor L862-0270 (10 µM) served as control. At 48 hpi, cells were fixed and ZsGreen-positive (ZsG^+^) signal was quantified using the Cytation 5 Microplate Reader (BioTek, Agilent) and normalized (%) to VC-treated infected controls. Results correspond to the mean and SD of four biological replicates. B) Representative IF images from panel A samples. Results were analyzed using Two-way ANOVA with multiple comparison correction with Šidák method. p value < 0.05, < 0.001 represented as ^*, ***^ respectively, while p value > 0.05 represented as ns (not significant).

### 3.3. Effect of cambinol on pH-dependent MaAv glycoprotein (GP)-mediated fusion

MaAv enter host cells through receptor-mediated endocytosis [49,50]. In the acidic environment of the endosome, subunit GP2 of the virus glycoprotein complex facilitates a pH-dependent membrane fusion between the viral and host membranes required to complete the cell entry process. To examine whether cambinol was also able to interfere with GP2-mediated fusion, we transfected HEK293T cells with plasmids encoding LCMV GPC (pC-LCMV-GPC), LASV GPC (pC-LASV-GPC), or an empty vector (pC-E) as a control, along with a plasmid expressing GFP (pC-GFP). At 5 h post-transfection, we treated cells with cambinol (100 μM) or VC. At 24 h post-transfection, we exposed cells to acidic (pH 5.0) or neutral (pH 7.2) medium for 15 min and then returned them to standard medium to monitor syncytia formation by GFP fluorescence (Figure 3). Cells expressing LCMV or LASV GPC exhibited robust syncytia formation under acidic conditions in the absence of cambinol, whereas cambinol treatment completely prevented syncytia formation, indicating that cambinol interferes with the GP2-mediated pH-dependent membrane fusion event required to complete the virus cell entry process.

**Figure 3.**
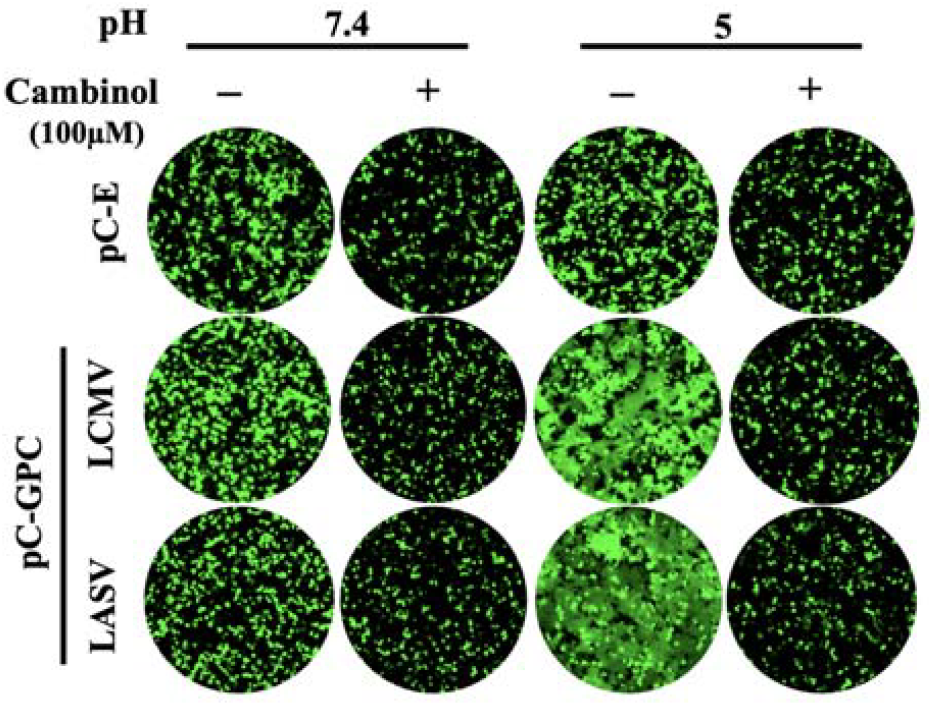
Effect of cambinol on LCMV and LASV GP-mediated fusion. HEK293T cells were seeded (5 × 10^5^ cells/well) onto poly-L-lysine-coated 24-well plates. At 16 h post-seeding, cells were transfected (4 µg/well) with plasmids encoding GPC from LCMV (pC-LCMV-GPC), or LASV (pC-LASV-GPC), or empty pCAGGS vector (pC-E) together with a GFP-expressing plasmid (pC-GFP; 50 ng/well). At 5 h post-transfection, cells were washed with DMEM supplemented with 10% FBS and treated with either cambinol (100 µM) or vehicle control (VC). At 24 h post-transfection, monolayers were exposed to acidified (pH 5.0) or neutral (pH 7.2) medium for 15 min and then returned to neutral medium (DMEM/10% FBS). The pattern of GFP expression was used to monitor fusion events over time, and once syncytia formation was apparent in VC-treated cells, samples were fixed (4% PFA) and IF images (4×) collected using a Keyence BZ-X710 microscope.

### 3.4. Effect of cambinol on LCMV vRNP Activity

As with other negative-strand RNA viruses, LCMV vRNP is responsible for directing the biosynthetic processes of replication and transcription of the viral genome. To evaluate the impact of cambinol on LCMV vRNP activity, we employed a cell-based LCMV minigenome (MG) system [24]. This system recapitulates LCMV RNA replication and transcription through intracellular reconstitution of the viral vRNP, which requires co-expression of the LCMV L and NP proteins together with a plasmid expressing the MG RNA. Levels of MG-directed GFP expression provide an integrated readout of vRNP activity. We transfected HEK293T cells in suspension with plasmids expressing the components of an LCMV MG system, and seeded them at 8 × 10^4^ cells/96-well. We used a MG system where the intracellularly reconstituted vRNP directed expression of GFP. Samples where the L-expressing plasmid was not included in the transfection mixture served as a negative control. Transfected cells were treated with cambinol (100 μM), vehicle control (VC), and VC containing 1%DMSO to match the DMSO concentration present in 100 μM cambinol generated from a 10 mM stock in 100% DMSO. At 72 h post-transfection, cells were fixed with 4% PFA and stained with DAPI. Representative IF images of each treatment condition (Figure 4A) were collected using a Keyence BZ-X710 microscope. GFP (Figure 4Bi) and DAPI (Figure 4Bii) signals were quantified using a Cytation 5 plate reader (BioTek, Agilent) and normalized (%) to VC-treated controls, which were set at 100% (Figure 4Ci,ii). Treatment with cambinol (100 μM) resulted in > 75% reduction in GFP expression compared to VC-treated controls. We did not observe a significant effect of cambinol on DAPI staining (Figure 4Cii), indicating that the modest cytostatic effect associated with cambinol treatment did not affect the activity of LCMV vRNP.

**Figure 4.**
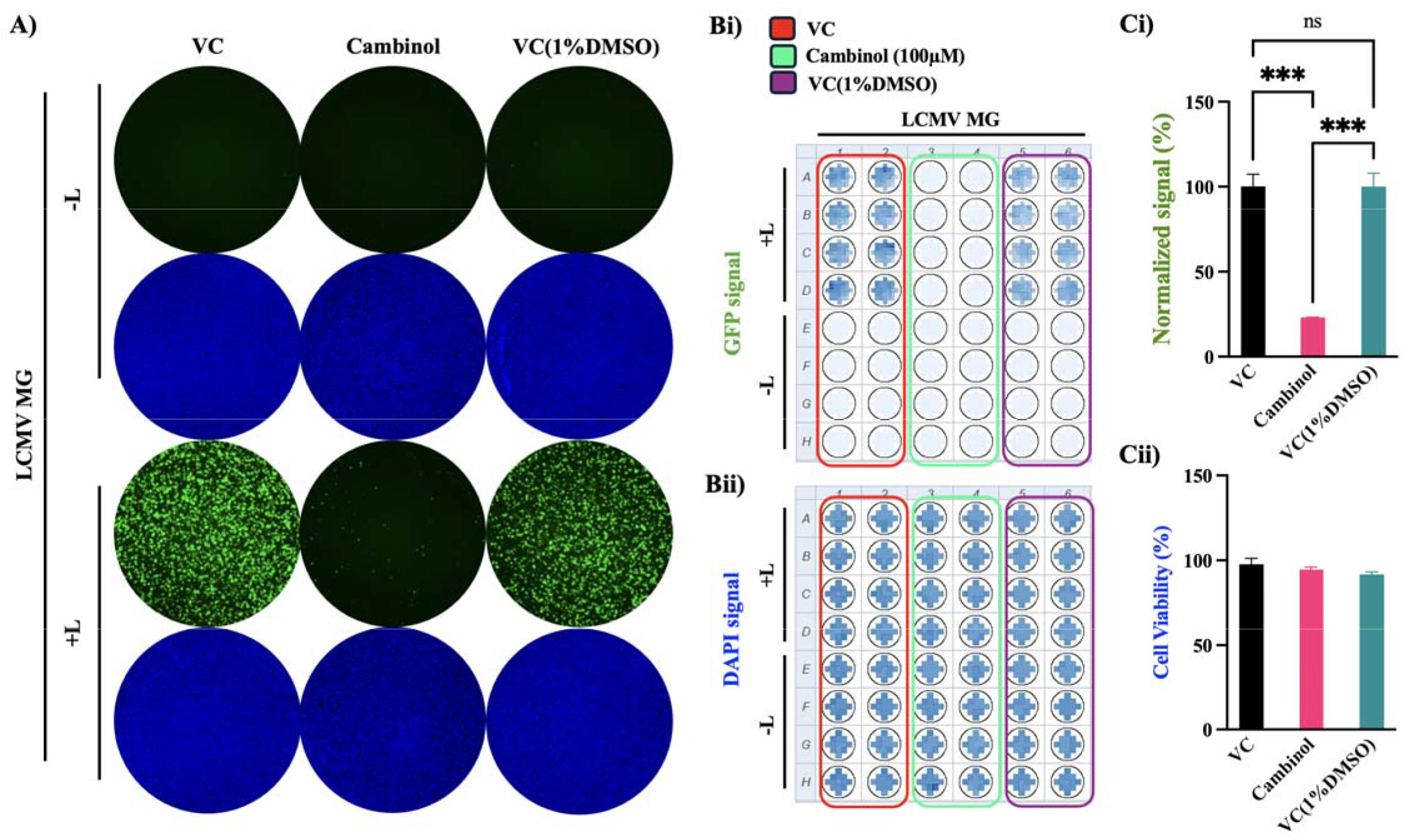
Effect of cambinol on LCMV vRNP activity. HEK293T cells were transfected in suspension with plasmids expressing the components of the LCMV T7-MG-GFP system (NP, L, and T7-MG-GFP) [24]. Samples where the L-expressing plasmid was not included in the transfection mixture served as a negative control for the MG activity. Transfected cells were seeded (80 µL/well containing 8 × 10^4^ cells) into 96-well plates pre-spotted with 20 µL of media containing either cambinol (100 µM), vehicle control (VC), and VC containing 1%DMSO, and mixed by pipetting twice to ensure homogenous distribution. At 72 h post-transfection, cells were fixed with 4% PFA and stained with DAPI. A) Representative IF images (4× magnification) from each treatment condition were collected using a Keyence BZ-X710 microscope. B) GFP (Bi) and DAPI (Bii) signals were quantified using a Cytation 5 plate reader (BioTek, Agilent). (C) GFP and DAPI signals were normalized (%) to VC-treated controls, which were set at 100% activity. Results represent the mean and SD of four independent replicates using seeding of cells at 6 × 10^4^ (Ci) or 8 × 10^4^ (Cii). D) Normalized GFP signal of samples treated with cambinol (100 µM) (Ci) and DAPI C(ii). Two-way ANOVA with multiple comparison correction with Dunnett method was implemented. p value < 0.001, represented as ^***^, while p value > 0.05 represented as ns (not significant).

### 3.5. Effect of cambinol on LCMV Z budding activity

The MaAv matrix Z protein is the main driver of virus budding [51,52]. To determine whether cambinol affected Z-mediated virus budding, we used an established cell-based Z budding assay where the activity of GLuc serves as a surrogate marker for Z budding activity [47]. We transfected HEK293T cells with a plasmid encoding a chimeric GLuc where its N-terminal secretory signal peptide was replaced with the Z open reading frame (Z-GLuc) of LCMV. Transfected cells were treated with cambinol (100 μM) or VC, and at 72 h post-transfection GLuc activity was measured in both CCSs containing virus-like particles (VLPs), and whole-cell lysates (WCLs). Z budding efficiency was calculated as the ratio of VLP-associated GLuc (GLuc in CCS) to total GLuc (GLuc CCS + GLuc-WCL) × 100. Values of budding efficiency were normalized (%) to those of VC-treated cells transfected with Z-WT that were assigned a value of 100%. As a control, we used Z(G2A)-GLuc, where mutation G2A prevents Z myristoylation, which results in inhibition of Z budding activity. Treatment with cambinol resulted in approximately 50% inhibition of Z-WT budding activity (Figure 5).

**Figure 5.**
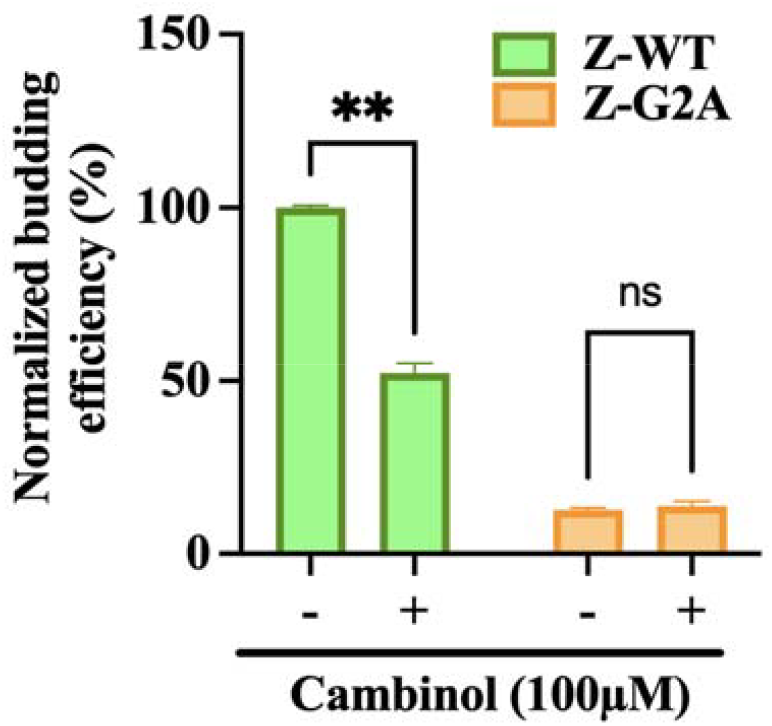
Effect of cambinol on LCMV Z budding activity. HEK293T cells were seeded onto poly-L-lysine-coated 12-well plate at a density of 5 × 10^5^ cells per well. Cells were transfected in suspension with plasmids encoding LCMV Z fused to Gaussia luciferase (GLuc) (pC-LCMV-Z-GLuc), or a myristoylation-deficient mutant form of Z (pC-LCMV-Z-G2A-GLuc) then seeded onto the plate. At 4 h post-transfection, media were replaced with fresh medium containing the indicated concentrations of the tested compounds or VC. At 72 h post-transfection, CCSs were collected and whole-cell lysates (WCL) prepared. GLuc activity in both CCS and WCL was quantified using the SteadyGlo Luciferase Pierce and measured with a Cytation 5 reader (BioTek, Agilent). GLuc activity in CCS served as a surrogate for Z-containing VLP released via Z budding activity, whereas GLuc activity in WCL reflected intracellular Z expression levels. Z budding efficiency was calculated as CCS-GLuc/(CCS-GLuc + WCL-GLuc) and normalized to VC-treated samples, set at 100%. Data were visualized using Prism 11. Average of technical triplicates is represented in the plot. Two-way ANOVA with Geisser-Greenhouse correction with Dunnett method was implemented. p value < 0.01, represented as ^**^, while p value > 0.05 represented as ns (not significant).

### 3.6. Contribution of Sirt-1 and Sirt-2 Isozymes to LCMV Propagation

To investigate a possible role of the two ubiquitously expressed sirtuin isozymes Sirt-1 and Sirt-2 on LCMV multiplication, we assessed multiplication of LCMV in HEK293 cells where expression of SIRT1 or SIRT2 was abrogated via CRISPR/Cas9 genetic editing as described [43], to generate SIRT1 KO and SIRT2 KO lines. We infected SIRT1 KO and SIRT2 KO cells with rLCMV/GFP (MOI = 0.05), and at the indicated hpi, we determined virus titers in CCS (Figure 6A). We fixed the same cell samples with 4% PFA and stained them with DAPI. GFP expression levels and DAPI staining signals were quantified using a Cytation 5 plate reader, and their values were normalized to those of VC-treated and infected cells. Normalized GFP and DAPI values were used to determine virus infectivity and cell viability, respectively (Figures 6B, C). Viral titers from the corresponding CCS samples (Figure 6A) were determined across independent biological replicates. Lack of Sirt-1 reduced production of infectious viral progeny by ∼ 1.2 logs at 48 hpi and by 0.8 log at 72 hpi (Figure 6A), which correlated with restricted virus propagation within the infected cell monolayer (Figure 6B, C). We observed in SIRT2 KO cells a ∼2.8-fold reduction at 48, but not at 72, hpi in infectious progeny production. Although this effect was rather modest, we cannot entirely exclude that Sirt-2 plays a transient supporting role early in the infection cycle, and future studies with Sirt-2 selective inhibitors would be required to unequivocally assess the contribution of Sirt-2 to LCMV, and in general MaAv, multiplication. These results suggest that Sirt-1 supports host cell functions required for optimal levels of LCMV multiplication, a finding consistent with the context-dependent pro-viral roles of Sirt-1 reported for other viruses [38].

**Figure 6.**
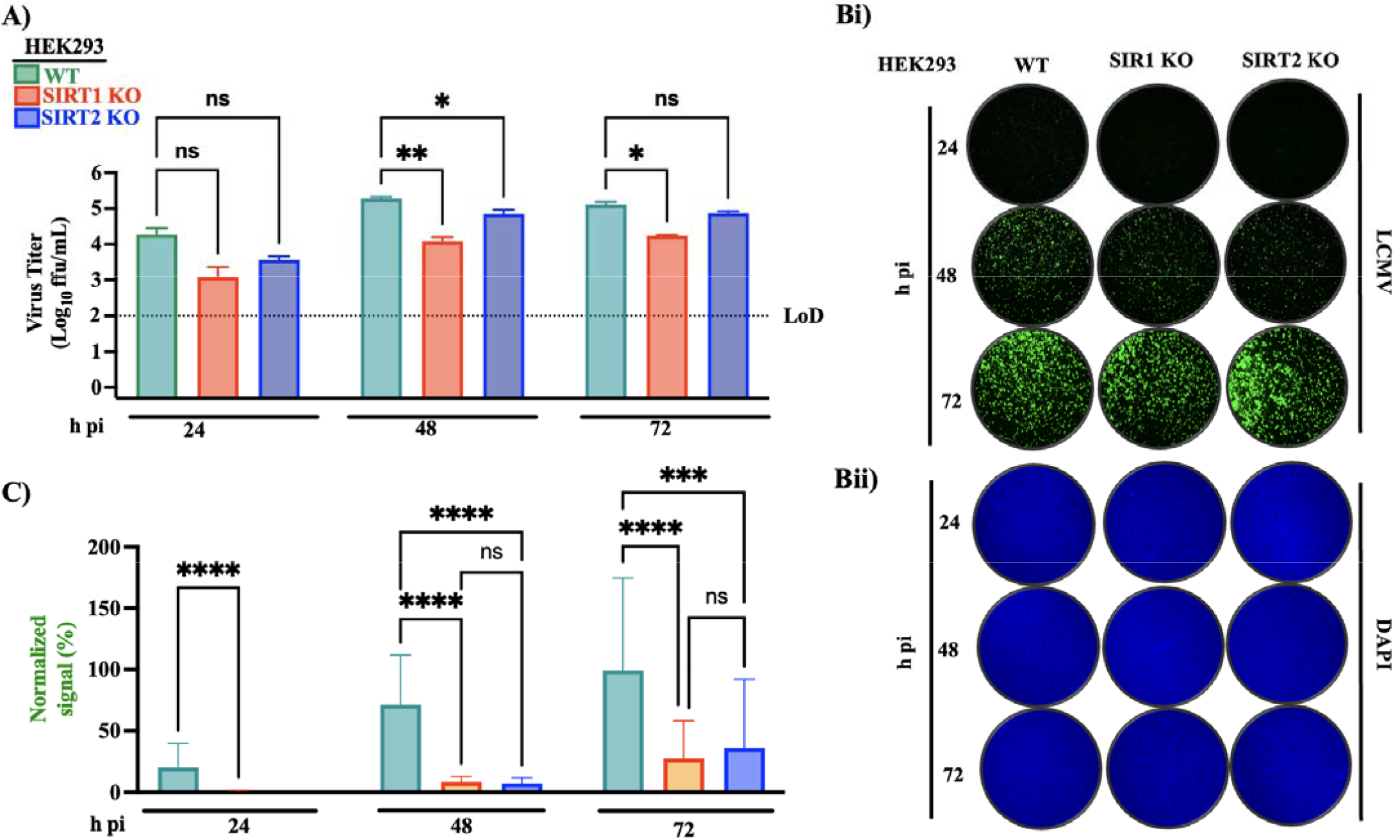
LCMV multiplication in SIRT1 and SIRT2 KO cells. A) HEK293 WT, SIRT1 or SIRT2 KO cells were seeded at 5 × 10^5^ cells/well into a 24-well plate, infected (MOI = 0.05) with rLCMV/GFP. At the indicated time points, CCSs were collected, and titers of infectious virus were determined by FFA using Vero E6 cells. B. At the indicated hpi, samples from (A) were fixed with 4% PFA and stained with DAPI (Bii). Images (4× magnification) were acquired using a Keyence BZ-X710 microscope. C. GFP signals were quantified using Cytation 5 and normalized to the average value of the VC-treated samples at 72 hpi. GFP and DAPI signals were quantified as the mean ± SD of 37 field scan measurements from a single well per condition, representing technical replicates within each well. Viral titers from the corresponding CCS samples (Figure 6A) were determined from technical triplicates. The two-way ANOVA analysis and Tukey’s correction for multiple comparisons were used. Statistically significant values: ns p > 0.05,^*^ p ≤ 0.05, ^**^ p < 0.01, ^***^ p < 0.001, ^****^ p < 0.0001.

### 3.7. Effect of SIRT1 or SIRT2 Knockout on LCMV cell entry and vRNP activity

To further investigate the mechanism by which SIRT1 or SIRT2 affected LCMV multiplication, we examined levels of ZsGreen expression in WT, SIRT1 and SIRT2 KO cells infected with the single-cycle infectious rLCMVΔGPC/ZsG and treated with the cell entry inhibitor F3406 at -1 hpi or +1 hpi, or with VC (Figure 7A). Treatment with F3406 starting at −1 hpi caused a similar drastic reduction in ZsGreen expression levels in all three cell lines (< 5% of the corresponding VC-treated sample). Treatment with F3406 starting at +1 hpi resulted in similar ZsGreen expression levels in all three cell lines (∼ 30% of those observed in the corresponding VC treated sample). These findings are consistent with SIRT1 and SIRT2 KO cell lines retaining a normal LCMV cell entry process. Under VC treatment conditions, SIRT1 KO, but not SIRT2 KO, cells exhibited a significant decrease in ZsGreen expression levels compared to those observed in WT cells, a finding supporting that LCMV vRNP-directed replication and transcription activities are impaired in SIRT1 KO cells, and consistent with LCMV multi-step growth kinetics results seen in WT, and SIRT1 and SIRT2 KO cells (Figure 6A).

**Figure 7.**
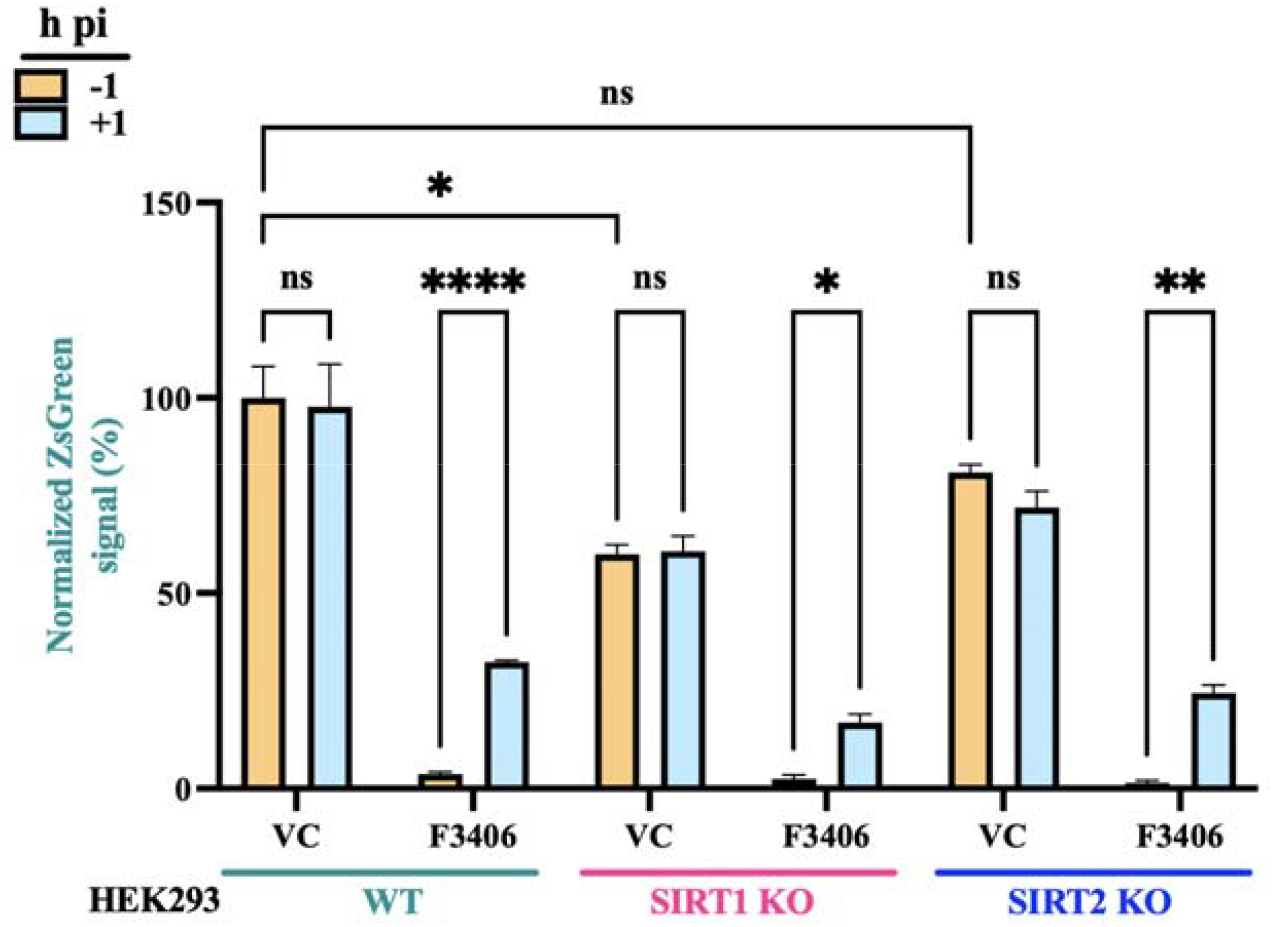
Effect of lack of SIRT1 or 2 on LCMV cell entry and vRNP activity. HEK293 WT, SIRT1 or SIRT2 KO cells were seeded in 96-well plates at 8 × 10^4^ cells/well. At 16 h post-seeding, cells were infected (MOI = 0.5) with a single-cycle infectious recombinant LCMV expressing ZsGreen (rLCMVΔGPC/ZsG). The LCMV entry inhibitor L862-0270 (10 µM) served as control. At 48 hpi, ZsGreen-positive (ZsG^+^) signal was quantified using the Cytation 5 Microplate Reader (BioTek, Agilent) and normalized (%) to VC-treated infected controls. Results correspond to the mean and SD of four biological replicates. Two-way ANOVA with multiple comparison correction with Šidák method was implemented. p value < 0.05, < 0.01, < 0.0001 represented as ^*, **, ****^ respectively, while p value > 0.05 represented as ns (not significant).

### 3.8. Effect of SIRT1 or SIRT2 Knockout on LCMV GP-Mediated Fusion

To examine whether SIRT1 or SIRT2 affected GP-mediated fusion, we transfected HEK293 WT, SIRT1 or 2 KO cells in suspension with plasmids encoding LCMV GPC (pC-LCMV-GPC), LASV GPC (pC-LASV-GPC), or an empty vector (pC-E) as a control, along with a plasmid expressing GFP (pC-GFP). Transfected cells were seeded into M24-well plates and at 24 h post-transfection exposed to acidic (pH 5.0) or neutral (pH 7.2) medium for 15 min and then returned to standard medium to monitor syncytia formation by GFP fluorescence (Figure 8). Upon exposure to low pH, LCMV or LASV GP exhibited robust fusion activity in WT, SIRT1 or 2 KO cells, indicating that lack of SIRT1 or 2 does not interfere with the pH-dependent GP2-mediated fusion activity.

**Figure 8.**
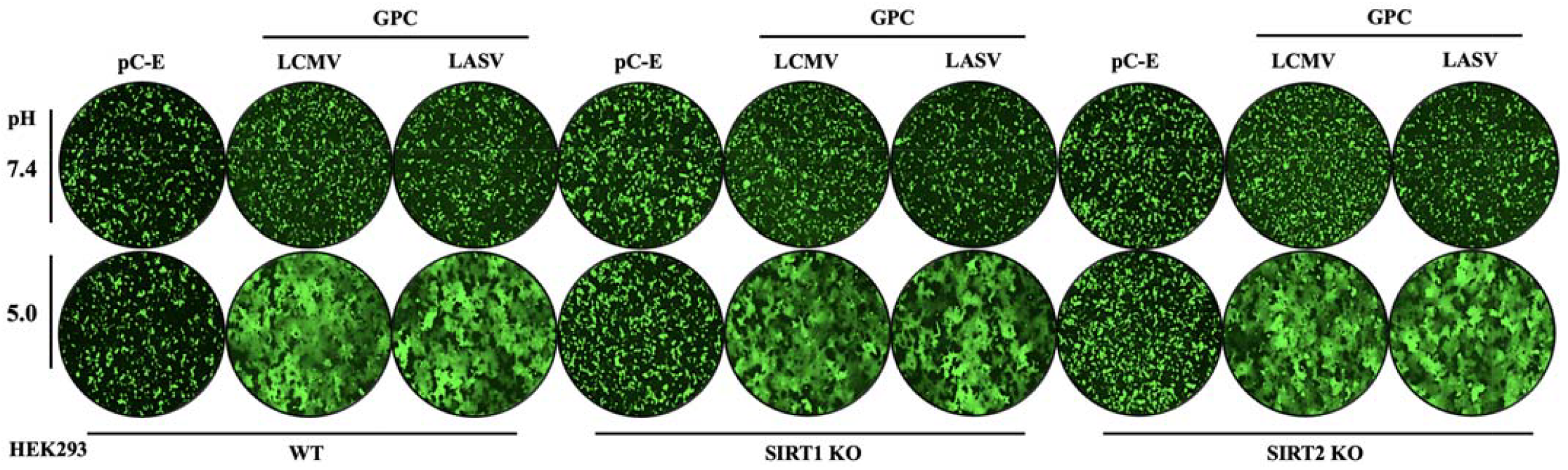
Effect of lack of SIRT1 or 2 on LCMV and LASV GPC-mediated fusion. HEK293 WT, SIRT1 or 2 KO cells were transferred (5 × 10^5^ cells/well) onto poly-L-lysine-coated 24-well plates and transfected immediately, then were set on a leveler for 2 minutes to settle down and establish attachment. The transfection mixture contained (1 µg/well) of plasmids encoding GPC from LCMV (pC-LCMV-GPC), or LASV (pC-LASV-GPC), or empty pCAGGS vector (pC-E) together with a GFP-expressing plasmid (pC-GFP; 50 ng/well). At 24 h post-transfection, monolayers were exposed to acidified (pH 5.0) or neutral (pH 7.2) medium for 15 min and then returned to neutral medium (DMEM/10% FBS). The pattern of GFP expression was used to monitor fusion events over time, and once syncytia formation was apparent in VC-treated cells, samples were fixed (4% PFA) and IF images (4×) collected using a Keyence BZ-X710 microscope.

### 3.9. Effect of SIRT1 or SIRT2 Knockout on Z budding activity

To examine whether Sirt-1 or Sirt-2 was required for Z-mediated budding, we transfected HEK293 WT, SIRT1 and SIRT2 KO cells with the Z-GLuc- or Z(G2A)-GLuc expressing plasmids of LCMV. At 72 h post-transfection, we measured GLuc activity in both CCSs, containing virus-like particles (VLPs), and whole-cell lysates (WCLs) and determined Z budding efficiency as described above in section 3.5 (Figure 9). Z budding efficiency was approximately five-fold higher in SIRT1 KO cells compared to wild-type cells, while SIRT2 KO cells showed no significant difference.

**Figure 9.**
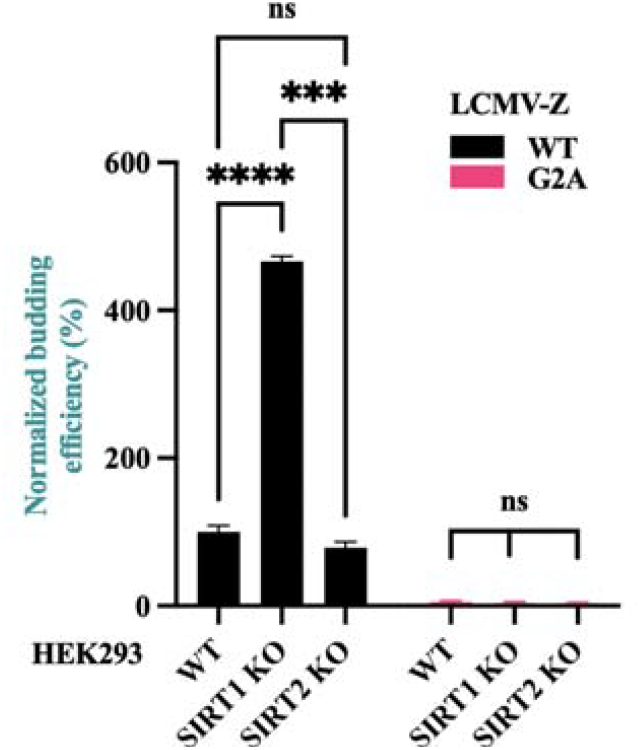
Effect of SIRT1 or SIRT2 Knockout on Z budding activity. HEK293 WT, SIRT1 or SIRT2 KO cells were seeded onto poly-L-lysine-coated 12-well plate at a density of 5 × 10^5^ cells per well. Cells were transfected in suspension with plasmids encoding LCMV Z fused to Gaussia luciferase (GLuc) (pC-LCMV-Z-GLuc), or a myristoylation-deficient mutant form of Z (pC-LCMV-Z-G2A-GLuc) then seeded onto the plate. At 72 h post-transfection, CCS were collected and whole-cell lysates (WCL) prepared. GLuc activity in both CCS and WCL was quantified using the SteadyGlo Luciferase Pierce and measured with a Cytation 5 reader (BioTek, Agilent). GLuc activity in CCS served as a surrogate for Z-containing VLP released via Z budding activity, while GLuc activity in WCL reflected intracellular Z expression levels. Z budding efficiency was calculated as CCS-GLuc/(CCS-GLuc + WCL-GLuc) and normalized to VC-treated samples, set at 100%. Data were visualized using Prism 11. Average of technical triplicates is represented in the plot. Two-way ANOVA with Geisser-Greenhouse correction with Dunnett method was implemented. p value < 0.001, < 0.0001 represented as ^***^,^****^ respectively, while p value > 0.05 represented as ns (not significant).

### 3.10. LCMV infection modulates SIRT pathway genes

Given the observed proviral role of SIRT in LCMV-infected cells, we sought to identify cellular genes modulated that SIRT that could contribute to the outcome of MaAv infection. For this, we re-analyzed a public dataset (GSE334699 available at Gene Expression Ominbus) containing RNAseq data from A549 cells infected with rLCMV/GFP [28]. We found 11 differentially expressed genes (DEGs) associated to SIRT pathway [53] (DOI: 10.1038/s41366-025-02007-w) in rLCMV/GFP infected cells meeting the criteria |log_2_ fold change infected/uninfected|> 2 and adjusted p value < 0.05 corrected by false discovery rate (FDR) (Figure 10). Of these, 10 genes were upregulated by infection (PARP8, PARP9, PARP10, PARP12, PARP14, PCK1, HK2, SOD22, DDIT3 and CD38) whereas only E2F1 was downregulated. These genes were related to important cellular pathways covering aspects of cellular metabolism, stress response, DNA-damage, FOXO mediated transcription, lipid metabolism or apoptosis among others. Overall, these results are consistent with LCMV modulation of SIRT pathway and further support the connection between SIRT and LCMV infection.

**Figure 10.**
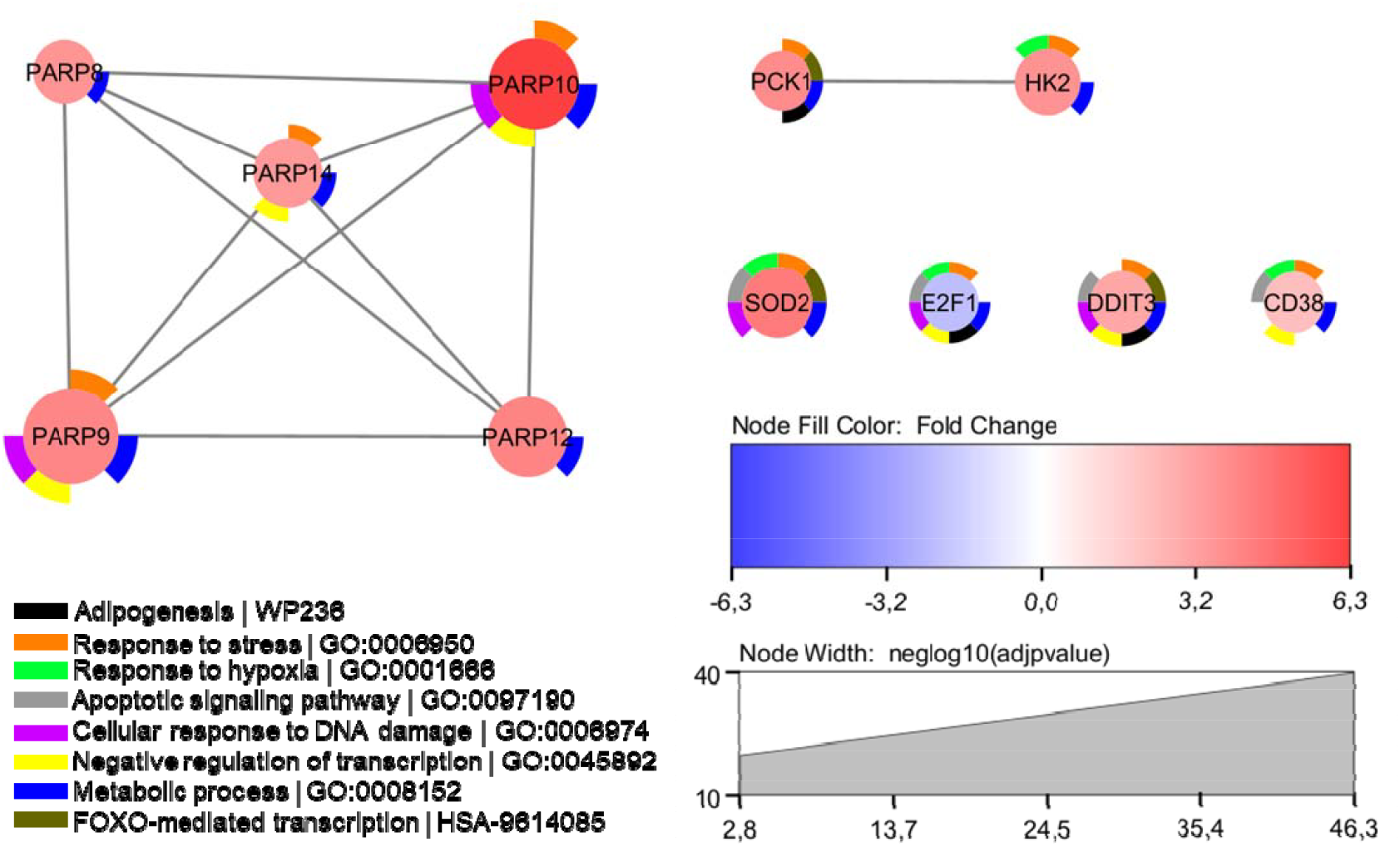
LCMV infection modulates SIRT pathway genes. SIRT pathway genes differenediatally expressed in A549 cells infected by rLCMV/GFP from GSE334699 dataset were selected on the basis of|log2 fold change infected/uninfected|> 2 and FDR-corrected P-value < 0.05. Donut slices indicate cellular pathways in which the genes are involved. Nodes were colored by fold change and node size denotes –log10 adjusted P-value. SIRT pathway genes were extracted from [53]. Network was created with STRING [54] and visualizad ysing Cytoscape [55].

## 4. Discussion

In this study, we show that cambinol inhibits several steps of the LCMV life cycle and identify Sirt-1 as a previously unrecognized pro-viral host factor that supports optimal LCMV multiplication. Our findings contribute to a growing body of evidence supporting an important role of host sphingolipid metabolism in MaAv multiplication.

We previously documented that cambinol exhibited a potent dose-dependent inhibitory effect on multiplication of rLCMV/GFP in Vero E6 cells [28]. Here we confirmed and expanded that observation by examining the dose-dependent effect of cambinol on rLCMV/GFP and an rLCMV/mCherry in human A549 cells (Figure 1). Both rLCMV/mCherry and rLCMV/GFP exhibited similar growth properties in A549 cells (Figure 1A) and cambinol inhibited both viruses in A549 with similar dose-dependent activity (Figure 1B, C). These results support the reproducibility of cambinol-mediated inhibition across reporter viruses and cell types.

Time-of-addition experiments demonstrated that cambinol targets both entry- and post-entry stages of LCMV life cycle. Consistent with this finding, cambinol inhibited the pH-dependent MaAv GP-mediated fusion event required for completion of MaAv cell entry, and vRNP-directed replication and transcription without significant effect on cell viability. Additionally, cambinol inhibited Z budding activity by approximately 50%, a finding consistent with the established dependence of MaAv Z-mediated VLP release on ceramide-enriched membrane microdomains [28–30,51,52,55–59].

We have shown that cambinol-mediated inhibition of nSMase2 results in inhibition of LCMV multiplication [28]. However, Sirt-1 and 2 are also validated targets of cambinol [60]. We therefore asked whether Sirt-1 and 2 contributed to the anti-LCMV activity of cambinol. LCMV multiplication was impaired in SIRT1, but not SIRT2, KO cells, revealing a previously unrecognized pro-viral role for Sirt-1 in the LCMV life cycle. This finding suggests that cambinol’s antiviral activity likely reflects the combined effects of targeting nSMase2 and Sirt-1.

Treatment with the LCMV entry inhibitor F3406 starting at -1 hpi, but not at +1 hpi, caused a very significant reduction in ZsGreen signal in SIRT1 and 2 KO cells infected with the single-cycle infectious rLCMVΔGPC/ZsG. In addition, pH-dependent GP-mediated fusion was not affected in SIRT1 or SIRT2 KO cells. These findings suggest that Sirt-1 primarily promotes post-cell entry events required for efficient LCMV multiplication.

Sirt-1 helps maintain lysosomal acidification through regulation of the vacuolar-type H^+^-ATPase proton pump [43,61,62]. Since LCMV entry requires trafficking through the endolysosomal compartment where acidic pH triggers the GP2-mediated fusion event necessary for vRNP release into the cytoplasm, impaired endolysosomal acidification in SIRT1 KO cells may negatively affect the efficiency of this step during LCMV infection. The reduced ZsGreen signal observed in SIRT1 KO cells, compared to SIRT2 KO and WT cells, in the time-of-addition assay may reflect contributions from both impaired endolysosomal acidification and vRNP activity (Figure 7). SIRT1 KO cells transfected with LCMV or LASV GPC exhibited syncytia formation upon exposure to low pH (Figure 8). However, in this assay cells over-expressing GPC are exposed to acidified medium allowing for the GP2-mediated fusion event to take place at the plasma membrane and bypassing the endolysosomal pathway.

Sirt-1 deacetylates a broad range of nuclear and cytoplasmic substrates involved in transcriptional regulation, RNA processing, stress responses, and autophagy [37]. Sirt-1-mediated regulation of the integrated stress response (ISR) and autophagy flux could influence the availability of cellular resources required for optimal vRNP function. Sirt-1 may also modulate the acetylation state of cellular RNA-binding proteins or translation factors that associate with the vRNP and contribute to its activity. Sirt-1 is predominantly nuclear, whereas LCMV replication and transcription take place in the cytoplasm. However, Sirt-1 is known to display nucleocytoplasmic shuttling. Whether the pro-viral effect of Sirt-1 reflects its direct cytoplasmic functions during LCMV infection, or its mediated effects on gene expression of host factors that support vRNP activity remains to be determined. Our results revealed that LCMV infection modulates the expression of several SIRT pathway related genes that are involved in multiple cellular processes including lipid metabolism and stress response, providing interesting candidates to further characterize the role of Sirt on MaAv infection. Acetylome profiling of infected SIRT1 KO and WT should help identify the specific substrates mediating the pro-viral activity of Sirt-1.

The significant (∼ five-fold) increase in Z budding activity observed in SIRT1 KO cells, despite markedly reduced overall viral multiplication, is consistent with the documented role of Sirt-1 as a negative regulator of production of extracellular vesicles (EV) [43], a process that is highly intertwined with the Z-mediated budding, which depends on interactions of Z canonical late-domain motifs with the ESCRT machinery [22,51,52,63–68]. Sirt-1 regulates nSMase2-dependent exosome biogenesis [43], and Sirt-1 loss may alter ceramide content and lipid raft organization in a manner that promotes Z-mediated VLP release. Consistent with this model, Sirt-1 deficiency has been shown to increase EV release in multiple cell systems by impairing lysosomal acidification and reducing degradation of multivesicular bodies [61,62]. SIRT1, but not SIRT2, KO cells also exhibit enlarged late endolysosomes and increased EV release [43]. These observations suggest that the increased Z-mediated VLP release in SIRT1 KO cells likely reflects a pro-secretory consequence of Sirt-1 deficiency on membrane budding pathways, rather than a virus-specific phenomenon. One possible explanation for the divergence between increased Z budding and reduced infectious output is that Sirt-1 loss uncouples particle release from productive genome replication or genome packaging. Impaired lysosomal acidification caused by reduced ATP6V1A expression can decrease multivesicular body degradation and increase fusion of multivesicular bodies with the plasma membrane [43,61,62]. Additionally, Sirt-1-dependent deacetylation of substrates involved in membrane trafficking or membrane scission could further enhance release of Z-driven particles [37]. This model parallels findings in HIV-1, where disruption of the coordination between genome packaging and budding results in non-infectious particle production [29,30]. Notably, compared to WT cells, SIRT2 KO cells showed no significant difference in Z budding efficiency, further supporting the isoform specificity of this regulatory effect and arguing that the pro-viral activity of Sirt-1 reflects its contribution to specific functions relevant to vRNP activity rather than a generic sirtuin class effect. The isoform specificity of the pro-viral effect is consistent with the distinct subcellular localizations and substrate repertoires of Sirt-1 and Sirt-2. Sirt-1 is predominantly nuclear and regulates transcriptional programs, stress responses, and autophagy, whereas Sirt-2 is primarily cytoplasmic with roles in cytoskeletal dynamics and mitotic regulation [37]. SIRT2 KO cells showed only a rather modest reduction in LCMV titers at 48 hpi that recovered to wild-type levels by 72 hpi, suggesting that under our experimental conditions Sirt-2 was not required for optimal LCMV multiplication. The selective effect of Sirt-1 on both vRNP activity and Z budding argues that the observed phenotypes do not reflect a generic consequence of sirtuin loss, but rather specific Sirt-1-regulated pathways that shape LCMV replication and particle release.

The pro-viral role of Sirt-1 in LCMV-infected cells contrasts with its reported antiviral activity in other virus-host systems including HCMV [38] and EV71 [39]. However, Sirt-1 can also promote viral replication as shown for HBV through transcription factor AP-1 [41] and for HIV-1 through Tat deacetylation [69]. Thus, the impact of Sirt-1 on viral infection is virus- and context-dependent, likely reflecting differences in viral replication strategy, host-cell type, and the specific Sirt-1-regulated pathways engaged during infection. The coordinated upregulation of metabolic enzymes (HK2, PCK1) and stress-response factors (DDIT3, CD38) suggests that LCMV exploits SIRT-dependent metabolic reprogramming to support viral replication. Exploration of sirtuin inhibitors as antiviral agents is warranted, though a comprehensive assessment of their therapeutic potential lies beyond the scope of the current study. Whether the pro-viral role identified here for LCMV extends to other MaAvs remains to be determined.

Several limitations of the present work should be considered. The experiments were conducted primarily in HEK293 cells, a transformed human kidney cell line that does not fully recapitulate the signaling environment of primary human target cells relevant to MaAv pathogenesis, including macrophages, hepatocytes, and endothelial cells. The molecular mechanisms by which Sirt-1 supports vRNP activity and modulates Z-mediated budding remain to be defined. It remains to be determined whether the increased Z budding efficiency in SIRT1 KO cells corresponds to increased total particle release with reduced genome incorporation or instead reflects altered membrane dynamics without proportional particle production. Electron microscopy, biochemical particle quantification, and genome-packaging analyses will be needed to distinguish among these possibilities. Future studies in primary human cells, organoid models, and animal models of MaAv infection will be required to establish the translational relevance of these findings and determine whether selective Sirt-1 inhibition could represent a viable host-directed antiviral strategy.

In summary, this study demonstrates that cambinol inhibits entry, vRNP activity, and Z-mediated budding of LCMV and identifies Sirt-1 as a novel pro-viral host factor required for optimal LCMV vRNP activity and infectious progeny production. These findings advance our understanding of the host pathways that govern MaAv multiplication and open new avenues for the development of host-directed antivirals against this important group of human pathogens.

## Author Contributions

Conceptualization, H.W., JCT; methodology, H.W.; validation, H.W.; formal analysis, R.K., M.A.M.A., P.C.M., H.W., JCT; investigation, H.W.; resources, H.W., JCT; writing R.K., H.W, P.C.M., A.B.B., M.A.M.A., JCT.; visualization, R.K. and H.W.; funding acquisition, M.A.M.A., H.W., JCT. All authors have read and agreed to the published version of the manuscript.

## Institutional Review Board Statement

Not applicable.

## Informed Consent Statement

Not applicable.

## Data Availability Statement

The original contributions presented in this study are included in the article/Supplementary Material. Further inquiries can be directed to the corresponding author.

## Acknowledgments

We thank Dr. Kwon (Medical College of Georgia, Augusta University) for providing us with the HEK293 sirtuin knockout cell lines.

## Abbreviations

The following abbreviations are used in this manuscript:

SIRT1: NAD^+^-dependent deacetylase sirtuin 1 gene
SIRT2: NAD^+^-dependent deacetylase sirtuin 2 gene
Sirt-1: NAD^+^-dependent deacetylase sirtuin 1
Sirt-2: NAD^+^-dependent deacetylase sirtuin 2

## References

1. Radoshitzky, S.R.; Buchmeier, M.; de la Torre, J.C. Emerging Viruses: Arenaviridae. In Fields Virology; Knipe, D. avid, Howley, P., Whelan, S., Eds.; 2020; Vol. I ISBN 978-1-9751-1254-7.

2. Basinski, A.J.; Fichet-Calvet, E.; Sjodin, A.R.; Varrelman, T.J.; Remien, C.H.; Layman, N.C.; Bird, B.H.; Wolking, D.J.; Monagin, C.; Ghersi, B.M.; et al. Bridging the Gap: Using Reservoir Ecology and Human Serosurveys to Estimate Lassa Virus Spillover in West Africa. PLOS Computational Biology 2021, 17, e1008811, doi:10.1371/journal.pcbi.1008811.

3. Fichet-Calvet, E.; Rogers, D.J. Risk Maps of Lassa Fever in West Africa. PLOS Neglected Tropical Diseases 2009, 3, e388, doi:10.1371/journal.pntd.0000388.

4. Grant, D.S.; Samuels, R.J.; Garry, R.F.; Schieffelin, J.S. Lassa Fever Natural History and Clinical Management. Curr Top Microbiol Immunol 2023, 440, 165–192, doi:10.1007/82_2023_263.

5. Sogoba, N.; Feldmann, H.; Safronetz, D. Lassa Fever in West Africa: Evidence for an Expanded Region of Endemicity. Zoonoses Public Health 2012, 59 Suppl 2, 43–47, doi:10.1111/j.1863-2378.2012.01469.x.

6. Klitting, R.; Kafetzopoulou, L.E.; Thiery, W.; Dudas, G.; Gryseels, S.; Kotamarthi, A.; Vrancken, B.; Gangavarapu, K.; Momoh, M.; Sandi, J.D.; et al. Predicting the Evolution of the Lassa Virus Endemic Area and Population at Risk over the next Decades. Nat Commun 2022, 13, 5596, doi:10.1038/s41467-022-33112-3.

7. Grant, A.; Seregin, A.; Huang, C.; Kolokoltsova, O.; Brasier, A.; Peters, C.; Paessler, S. Junín Virus Pathogenesis and Virus Replication. Viruses 2012, 4, 2317–2339, doi:10.3390/v4102317.

8. Lendino, A.; Castellanos, A.A.; Pigott, D.M.; Han, B.A. A Review of Emerging Health Threats from Zoonotic New World Mammarenaviruses. BMC Microbiology 2024, 24, 115, doi:10.1186/s12866-024-03257-w.

9. Bonthius, D.J. Lymphocytic Choriomeningitis Virus: A Prenatal and Postnatal Threat. Adv Pediatr 2009, 56, 75–86, doi:10.1016/j.yapd.2009.08.007.

10. Palacios, G.; Druce, J.; Du, L.; Tran, T.; Birch, C.; Briese, T.; Conlan, S.; Quan, P.-L.; Hui, J.; Marshall, J.; et al. A New Arenavirus in a Cluster of Fatal Transplant-Associated Diseases. N Engl J Med 2008, 358, 991–998, doi:10.1056/nejmoa073785.

11. Bonthius, D.J. Lymphocytic Choriomeningitis Virus: An Underrecognized Cause of Neurologic Disease in the Fetus, Child, and Adult. Semin Pediatr Neurol 2012, 19, 89–95, doi:10.1016/j.spen.2012.02.002.

12. Vilibic-Cavlek, T.; Savic, V.; Ferenc, T.; Mrzljak, A.; Barbic, L.; Bogdanic, M.; Stevanovic, V.; Tabain, I.; Ferencak, I.; Zidovec-Lepej, S. Lymphocytic Choriomeningitis—Emerging Trends of a Neglected Virus: A Narrative Review. Trop Med Infect Dis 2021, 6, 88, doi:10.3390/tropicalmed6020088.

13. Salam, A.P.; Cheng, V.; Edwards, T.; Olliaro, P.; Sterne, J.; Horby, P. Time to Reconsider the Role of Ribavirin in Lassa Fever. PLOS Neglected Tropical Diseases 2021, 15, e0009522, doi:10.1371/journal.pntd.0009522.

14. Mendenhall, M.; Russell, A.; Juelich, T.; Messina, E.L.; Smee, D.F.; Freiberg, A.N.; Holbrook, M.R.; Furuta, Y.; de la Torre, J.-C.; Nunberg, J.H.; et al. T-705 (Favipiravir) Inhibition of Arenavirus Replication in Cell Culture. Antimicrob Agents Chemother 2011, 55, 782–787, doi:10.1128/AAC.01219-10.

15. Welch, S.R.; Spengler, J.R.; Westover, J.B.; Bailey, K.W.; Davies, K.A.; Aida-Ficken, V.; Bluemling, G.R.; Boardman, K.M.; Wasson, S.R.; Mao, S.; et al. Delayed Low-Dose Oral Administration of 4′-Fluorouridine Inhibits Pathogenic Arenaviruses in Animal Models of Lethal Disease. Science Translational Medicine 2024, 16, eado7034, doi:10.1126/scitranslmed.ado7034.

16. Cashman, K.A.; Wilkinson, E.R.; Posakony, J.; Madu, I.G.; Tarcha, E.J.; Lustig, K.H.; Korth, M.J.; Bedard, K.M.; Amberg, S.M. Lassa Antiviral LHF-535 Protects Guinea Pigs from Lethal Challenge. Sci Rep 2022, 12, 19911, doi:10.1038/s41598-022-23760-2.

17. Gowen, B.B.; Naik, S.; Westover, J.B.; Brown, E.R.; Gantla, V.R.; Fetsko, A.; Dagley, A.L.; Blotter, D.J.; Anderson, N.; McCormack, K.; et al. Potent Inhibition of Arenavirus Infection by a Novel Fusion Inhibitor. Antiviral Res 2021, 193, 105125, doi:10.1016/j.antiviral.2021.105125.

18. Cross, R.W.; Hastie, K.M.; Mire, C.E.; Robinson, J.E.; Geisbert, T.W.; Branco, L.M.; Ollmann Saphire, E.; Garry, R.F. Antibody Therapy for Lassa Fever. Curr Opin Virol 2019, 37, 97–104, doi:10.1016/j.coviro.2019.07.003.

19. Eudy, E.; Woodburn, D.; Reeder, R.; Cooper, K.; Hart, R.; Hischak, A.; Eaton, B.; Ruedas, J.B.; Tran, T.; Johnson, C.; et al. The Virus Entry Inhibitor ARN-75039 Provides Therapeutic Protection against Lassa Virus Infection in Guinea Pigs. Science Translational Medicine 2026, 18, eadx0938, doi:10.1126/scitranslmed.adx0938.

20. Erameh, C.; Okwaraeke, K.; Kleist, C.; Edeawe, O.; Adedosu, N.; Ekata, E.; Abejegah, C.; Owhin, S.; Babatunde, F.; Omansen, T.; et al. Favipiravir for Lassa Fever: An Open-Label, Randomized Controlled Phase 2 Trial. Nat Med 2026, 1–11, doi:10.1038/s41591-026-04402-w.

21. Kumar, N.; Sharma, S.; Kumar, R.; Tripathi, B.N.; Barua, S.; Ly, H.; Rouse, B.T. Host-Directed Antiviral Therapy. Clin Microbiol Rev 2020, 33, e00168–19, doi:10.1128/CMR.00168-19.

22. Witwit, H.; Betancourt, C.A.; Cubitt, B.; Khafaji, R.; Kowalski, H.; Jackson, N.; Ye, C.; Martinez-Sobrido, L.; de la Torre, J.C. Cellular N-Myristoyl Transferases Are Required for Mammarenavirus Multiplication. Viruses 2024, 16, 1362, doi:10.3390/v16091362.

23. Witwit, H.; de la Torre, J.C. N-Myristoyltransferase Inhibitors as Candidate Broad-Spectrum Antivirals to Treat Viral Infections Promoted by Immunosuppression Associated with JAK Inhibitors Therapy. Antiviral Research 2025, 242, 106258, doi:10.1016/j.antiviral.2025.106258.

24. Witwit, H.; Ibanez, P.A.; Zhou, R.; Jackson, N.; Escobedo, R.; Cubitt, B.; Khafaji, R.; Sattler, R.Y.; Martinez-Sobrido, L.; de la Torre, J.C. Prolyl tRNA Synthetase Is Required for Mammarenavirus Multiplication. Viruses 2026, 18, 202, doi:10.3390/v18020202.

25. Kim, Y.-J.; Witwit, H.; Cubitt, B.; de la Torre, J.C. Inhibitors of Anti-Apoptotic Bcl-2 Family Proteins Exhibit Potent and Broad-Spectrum Anti-Mammarenavirus Activity via Cell Cycle Arrest at G0/G1 Phase. Journal of Virology 2021, 95, 10.1128/jvi.01399-21, doi:10.1128/jvi.01399-21.

26. Zumla, A.; Rao, M.; Wallis, R.S.; Kaufmann, S.H.E.; Rustomjee, R.; Mwaba, P.; Vilaplana, C.; Yeboah-Manu, D.; Chakaya, J.; Ippolito, G.; et al. Host-Directed Therapies for Infectious Diseases: Current Status, Recent Progress, and Future Prospects. Lancet Infect Dis 2016, 16, e47–63, doi:10.1016/S1473-3099(16)00078-5.

27. Han, S.; Ye, X.; Yang, J.; Peng, X.; Jiang, X.; Li, J.; Zheng, X.; Zhang, X.; Zhang, Y.; Zhang, L.; et al. Host Specific Sphingomyelin Is Critical for Replication of Diverse RNA Viruses. Cell Chem Biol 2024, 31, 2052–2068.e11, doi:10.1016/j.chembiol.2024.10.009.

28. Mingo-Casas, P.; Witwit, H.; Casasampere, M.; Blazquez, A.B.; Cubitt, B.; Martin-Acebes, M.A.; Torre, J.C. de la Mammarenavirus-Induced Remodeling of the Cellular Lipid Landscape Reveals Sphingolipid Metabolism as a Novel Target for Antiviral Intervention 2026, 2026.07.02.736094.

29. Yoo, S.-W.; Waheed, A.A.; Deme, P.; Tohumeken, S.; Rais, R.; Smith, M.D.; DeMarino, C.; Calabresi, P.A.; Kashanchi, F.; Freed, E.O.; et al. Inhibition of Neutral Sphingomyelinase 2 Impairs HIV-1 Envelope Formation and Substantially Delays or Eliminates Viral Rebound. Proceedings of the National Academy of Sciences 2023, 120, e2219543120, doi:10.1073/pnas.2219543120.

30. Waheed, A.A.; Zhu, Y.; Agostino, E.; Naing, L.; Hikichi, Y.; Soheilian, F.; Yoo, S.-W.; Song, Y.; Zhang, P.; Slusher, B.S.; et al. Neutral Sphingomyelinase 2 Is Required for HIV-1 Maturation. Proceedings of the National Academy of Sciences 2023, 120, e2219475120, doi:10.1073/pnas.2219475120.

31. Heltweg, B.; Gatbonton, T.; Schuler, A.D.; Posakony, J.; Li, H.; Goehle, S.; Kollipara, R.; Depinho, R.A.; Gu, Y.; Simon, J.A.; et al. Antitumor Activity of a Small-Molecule Inhibitor of Human Silent Information Regulator 2 Enzymes. Cancer Res 2006, 66, 4368–4377, doi:10.1158/0008-5472.CAN-05-3617.

32. Airola, M.V.; Hannun, Y.A. Sphingolipid Metabolism and Neutral Sphingomyelinases. Handb Exp Pharmacol 2013, 57–76, doi:10.1007/978-3-7091-1368-4_3.

33. Menck, K.; Sönmezer, C.; Worst, T.S.; Schulz, M.; Dihazi, G.H.; Streit, F.; Erdmann, G.; Kling, S.; Boutros, M.; Binder, C.; et al. Neutral Sphingomyelinases Control Extracellular Vesicles Budding from the Plasma Membrane. J Extracell Vesicles 2017, 6, 1378056, doi:10.1080/20013078.2017.1378056.

34. Back, M.J.; Ha, H.C.; Fu, Z.; Choi, J.M.; Piao, Y.; Won, J.H.; Jang, J.M.; Shin, I.C.; Kim, D.K. Activation of Neutral Sphingomyelinase 2 by Starvation Induces Cell-Protective Autophagy via an Increase in Golgi-Localized Ceramide. Cell Death Dis 2018, 9, 670, doi:10.1038/s41419-018-0709-4.

35. Álvarez-Fernández, H.; Mingo-Casas, P.; Blázquez, A.-B.; Caridi, F.; Saiz, J.C.; Pérez-Pérez, M.-J.; Martín-Acebes, M.A.; Priego, E.-M. Allosteric Inhibition of Neutral Sphingomyelinase 2 (nSMase2) by DPTIP: From Antiflaviviral Activity to Deciphering Its Binding Site through In Silico Studies and Experimental Validation. Int J Mol Sci 2022, 23, 13935, doi:10.3390/ijms232213935.

36. Salisch, F.; Schumacher, F.; Gärtner, U.; Kleuser, B.; Ziebuhr, J.; Müller-Ruttloff, C. Targeting Sphingolipid Metabolism: Inhibition of Neutral Sphingomyelinase 2 Impairs Coronaviral Replication Organelle Formation. mBio 2025, 16, e00084–25, doi:10.1128/mbio.00084-25.

37. Chang, H.-C.; Guarente, L. SIRT1 and Other Sirtuins in Metabolism. Trends Endocrinol Metab 2014, 25, 138–145, doi:10.1016/j.tem.2013.12.001.

38. Koyuncu, E.; Budayeva, H.G.; Miteva, Y.V.; Ricci, D.P.; Silhavy, T.J.; Shenk, T.; Cristea, I.M. Sirtuins Are Evolutionarily Conserved Viral Restriction Factors. mBio 2014, 5, e02249–14, doi:10.1128/mBio.02249-14.

39. Han, Y.; Wang, L.; Cui, J.; Song, Y.; Luo, Z.; Chen, J.; Xiong, Y.; Zhang, Q.; Liu, F.; Ho, W.; et al. SIRT1 Inhibits EV71 Genome Replication and RNA Translation by Interfering with the Viral Polymerase and 5′UTR RNA. J Cell Sci 2016, 129, 4534–4547, doi:10.1242/jcs.193698.

40. He, M.; Gao, S.-J. A Novel Role of SIRT1 in Gammaherpesvirus Latency and Replication. Cell Cycle 2014, 13, 3328–3330, doi:10.4161/15384101.2014.968431.

41. Ren, J.-H.; Tao, Y.; Zhang, Z.-Z.; Chen, W.-X.; Cai, X.-F.; Chen, K.; Ko, B.C.B.; Song, C.-L.; Ran, L.-K.; Li, W.-Y.; et al. Sirtuin 1 Regulates Hepatitis B Virus Transcription and Replication by Targeting Transcription Factor AP-1. J Virol 2014, 88, 2442–2451, doi:10.1128/JVI.02861-13.

42. Süssmuth, S.D.; Haider, S.; Landwehrmeyer, G.B.; Farmer, R.; Frost, C.; Tripepi, G.; Andersen, C.A.; Di Bacco, M.; Lamanna, C.; Diodato, E.; et al. An Exploratory Double-Blind, Randomized Clinical Trial with Selisistat, a SirT1 Inhibitor, in Patients with Huntington’s Disease. Br J Clin Pharmacol 2015, 79, 465–476, doi:10.1111/bcp.12512.

43. Lee, B.R.; Sanstrum, B.J.; Liu, Y.; Kwon, S.-H. Distinct Role of Sirtuin 1 (SIRT1) and Sirtuin 2 (SIRT2) in Inhibiting Cargo-Loading and Release of Extracellular Vesicles. Sci Rep 2019, 9, 20049, doi:10.1038/s41598-019-56635-0.

44. Iwasaki, M.; Minder, P.; Caì, Y.; Kuhn, J.H.; Yates, J.R.; Torbett, B.E.; de la Torre, J.C. Interactome Analysis of the Lymphocytic Choriomeningitis Virus Nucleoprotein in Infected Cells Reveals ATPase Na+/K+ Transporting Subunit Alpha 1 and Prohibitin as Host-Cell Factors Involved in the Life Cycle of Mammarenaviruses. PLoS Pathog 2018, 14, e1006892, doi:10.1371/journal.ppat.1006892.

45. Sánchez, A.B.; de la Torre, J.C. Rescue of the Prototypic Arenavirus LCMV Entirely from Plasmid. Virology 2006, 350, 370–380, doi:10.1016/j.virol.2006.01.012.

46. Battegay, M.; Cooper, S.; Althage, A.; Bänziger, J.; Hengartner, H.; Zinkernagel, R.M. Quantification of Lymphocytic Choriomeningitis Virus with an Immunological Focus Assay in 24-or 96-Well Plates. J Virol Methods 1991, 33, 191–198, doi:10.1016/0166-0934(91)90018-u.

47. Capul, A.A.; de la Torre, J.C. A Cell-Based Luciferase Assay Amenable to High-Throughput Screening of Inhibitors of Arenavirus Budding. Virology 2008, 382, 107–114, doi:10.1016/j.virol.2008.09.008.

48. Quirin, K.; Eschli, B.; Scheu, I.; Poort, L.; Kartenbeck, J.; Helenius, A. Lymphocytic Choriomeningitis Virus Uses a Novel Endocytic Pathway for Infectious Entry via Late Endosomes. Virology 2008, 378, 21–33, doi:10.1016/j.virol.2008.04.046.

49. Borrow, P.; Oldstone, M.B. Mechanism of Lymphocytic Choriomeningitis Virus Entry into Cells. Virology 1994, 198, 1–9, doi:10.1006/viro.1994.1001.

50. White, J.; Helenius, A. pH-Dependent Fusion between the Semliki Forest Virus Membrane and Liposomes. Proc Natl Acad Sci U S A 1980, 77, 3273–3277, doi:10.1073/pnas.77.6.3273.

51. Perez, M.; Greenwald, D.L.; de La Torre, J.C. Myristoylation of the RING Finger Z Protein Is Essential for Arenavirus Budding. Journal of Virology 2004, 78, 11443–11448, doi:10.1128/jvi.78.20.11443-11448.2004.

52. Witwit, H.; de la Torre, J.C. Mammarenavirus Z Protein Myristoylation and Oligomerization Are Not Required for Its Dose-Dependent Inhibitory Effect on vRNP Activity. BioChem 2025, 5, 10, doi:10.3390/biochem5020010.

53. Szklarczyk, D.; Kirsch, R.; Koutrouli, M.; Nastou, K.; Mehryary, F.; Hachilif, R.; Gable, A.L.; Fang, T.; Doncheva, N.T.; Pyysalo, S.; et al. The STRING Database in 2023: Protein-Protein Association Networks and Functional Enrichment Analyses for Any Sequenced Genome of Interest. Nucleic Acids Res 2023, 51, D638–D646, doi:10.1093/nar/gkac1000.

54. Shannon, P.; Markiel, A.; Ozier, O.; Baliga, N.S.; Wang, J.T.; Ramage, D.; Amin, N.; Schwikowski, B.; Ideker, T. Cytoscape: A Software Environment for Integrated Models of Biomolecular Interaction Networks. Genome Research 2003, 13, 2498–2504, doi:10.1101/gr.1239303.

55. Seo, Y.-J.; Hahm, B. Sphingosine Analog AAL-R Promotes Activation of LCMV-Infected Dendritic Cells. Viral Immunol 2014, 27, 82–86, doi:10.1089/vim.2013.0096.

56. Pritzl, C.J.; Seo, Y.-J.; Xia, C.; Vijayan, M.; Stokes, Z.D.; Hahm, B. A Ceramide Analogue Stimulates Dendritic Cells to Promote T Cell Responses upon Virus Infections. J Immunol 2015, 194, 4339–4349, doi:10.4049/jimmunol.1402672.

57. Hu, Z.; Abdelrahman, H.; Elwy, A.; Kuang, F.; Dhiman, S.; Ali, S.; Marson, M.; Holnsteiner, L.; Hamdan, T.A.; Friebus-Kardash, J.; et al. Acid Ceramidase Regulates CD8+ T-Cell Exhaustion via Type I Interferon-Mediated Upregulation of PD-L1. Front. Immunol. 2025, 16, doi:10.3389/fimmu.2025.1638403.

58. Hose, M.; Günther, A.; Naser, E.; Schumacher, F.; Schönberger, T.; Falkenstein, J.; Papadamakis, A.; Kleuser, B.; Becker, K.A.; Gulbins, E.; et al. Cell-Intrinsic Ceramides Determine T Cell Function during Melanoma Progression. eLife 2022, 11, e83073, doi:10.7554/eLife.83073.

59. Avota, E.; Bodem, J.; Chithelen, J.; Mandasari, P.; Beyersdorf, N.; Schneider-Schaulies, J. The Manifold Roles of Sphingolipids in Viral Infections. Front Physiol 2021, 12, 715527, doi:10.3389/fphys.2021.715527.

60. Giordano, D.; Scafuri, B.; De Masi, L.; Capasso, L.; Maresca, V.; Altucci, L.; Nebbioso, A.; Facchiano, A.; Bontempo, P. Sirtuin Inhibitor Cambinol Induces Cell Differentiation and Differently Interferes with SIRT1 and 2 at the Substrate Binding Site. Biomedicines 2023, 11, 1624, doi:10.3390/biomedicines11061624.

61. Ding, L.; Li, Z.; Zhou, Y.; Liu, N.; Liu, S.; Zhang, X.; Liu, C.; Zhang, D.; Wang, G.; Ma, R. Loss of Sirt1 Promotes Exosome Secretion from Podocytes by Inhibiting Lysosomal Acidification in Diabetic Nephropathy. Molecular and Cellular Endocrinology 2023, 568–569, 111913, doi:10.1016/j.mce.2023.111913.

62. Zhang, S.; Yang, Y.; Lv, X.; Zhou, X.; Zhao, W.; Meng, L.; Xu, H.; Zhu, S.; Wang, Y. Doxorubicin-Induced Cardiotoxicity Through SIRT1 Loss Potentiates Overproduction of Exosomes in Cardiomyocytes. Int J Mol Sci 2024, 25, 12376, doi:10.3390/ijms252212376.

63. Strecker, T.; Maisa, A.; Daffis, S.; Eichler, R.; Lenz, O.; Garten, W. The Role of Myristoylation in the Membrane Association of the Lassa Virus Matrix Protein Z. Virol J 2006, 3, 93, doi:10.1186/1743-422X-3-93.

64. Strecker, T.; Eichler, R.; Meulen, J. ter; Weissenhorn, W.; Dieter Klenk, H.; Garten, W.; Lenz, O. Lassa Virus Z Protein Is a Matrix Protein Sufficient for the Release of Virus-Like Particles. J Virol 2003, 77, 10700–10705, doi:10.1128/JVI.77.19.10700-10705.2003.

65. Cornu, T.I.; de la Torre, J.C. RING Finger Z Protein of Lymphocytic Choriomeningitis Virus (LCMV) Inhibits Transcription and RNA Replication of an LCMV S-Segment Minigenome. J Virol 2001, 75, 9415–9426, doi:10.1128/JVI.75.19.9415-9426.2001.

66. Liu, L.; Wang, P.; Liu, A.; Zhang, L.; Yan, L.; Guo, Y.; Xiao, G.; Rao, Z.; Lou, Z. Structure Basis for Allosteric Regulation of Lymphocytic Choriomeningitis Virus Polymerase Function by Z Matrix Protein. protein. cell. 2023, 14, 703–707, doi:10.1093/procel/pwad018.

67. Ziegler, C.M.; Dang, L.; Eisenhauer, P.; Kelly, J.A.; King, B.R.; Klaus, J.P.; Manuelyan, I.; Mattice, E.B.; Shirley, D.J.; Weir, M.E.; et al. NEDD4 Family Ubiquitin Ligases Associate with LCMV Z’s PPXY Domain and Are Required for Virus Budding, but Not via Direct Ubiquitination of Z. PLoS Pathog 2019, 15, e1008100, doi:10.1371/journal.ppat.1008100.

68. Kranzusch, P.J.; Whelan, S.P.J. Arenavirus Z Protein Controls Viral RNA Synthesis by Locking a Polymerase–Promoter Complex. Proceedings of the National Academy of Sciences 2011, 108, 19743–19748, doi:10.1073/pnas.1112742108.

69. Pagans, S.; Pedal, A.; North, B.J.; Kaehlcke, K.; Marshall, B.L.; Dorr, A.; Hetzer-Egger, C.; Henklein, P.; Frye, R.; McBurney, M.W.; et al. SIRT1 Regulates HIV Transcription via Tat Deacetylation. PLOS Biology 2005, 3, e41, doi:10.1371/journal.pbio.0030041.

